# A Multi-Omic Atlas of Convergent and Divergent Metabolic Regulatory Circuitries in Cancer

**DOI:** 10.1101/2025.11.15.688631

**Authors:** Higor Almeida Cordeiro Nogueira, Emanuell Rodrigues de Souza, Victor dos Santos Lopes, Enrique Medina-Acosta

**Affiliations:** Laboratório de Biotecnologia, Centro de Biociências e Biotecnologia, Universidade Estadual do Norte Fluminense, Campos dos Goytacazes, Brazil

**Keywords:** cancer metabolism, metabolic reprogramming, metabolic regulatory circuitries, omic-specific metabolic signatures, OncoMetabolismGPS, regulated cell death

## Abstract

Metabolic reprogramming underlies tumor progression, immune evasion, and resistance to regulated cell death, yet the higher-order regulatory logic that coordinates these processes across molecular layers remains poorly defined. We developed OncoMetabolismGPS, a multi-omic analytical framework that reconstructs a Pan-Cancer atlas of convergent and divergent metabolic regulatory circuitries. From 463,433 significant multi-omic and phenotypic associations across 33 tumor types, we derived 241,415 omic-specific metabolic signatures, each integrating metabolic pathway context with phenotypic, prognostic, and immune features. By mapping shared upstream regulators of these signatures, we identified 24,796 metabolic regulatory circuitries—classified as convergent when regulators and signatures act in the same biological direction, or divergent when they exhibit opposing associations. Divergent circuitry predominated, especially in immunosuppressive (cold) tumor contexts, revealing context-dependent regulatory compensation across metabolic, phenotypic, and clinical axes. The accompanying OncoMetabolismGPS Shiny application implements this atlas as an interactive platform that positions each signature and circuitry within a multidimensional coordinate space defined by molecular, phenotypic, immune, and clinical attributes, enabling systematic navigation of metabolic regulatory behavior in cancer. Together, this study establishes the first multi-omic atlas of metabolic regulatory circuitries, providing a conceptual and computational framework for dissecting metabolic plasticity, pathway dependencies, and therapeutic vulnerabilities across human cancers.

## 1. Introduction

The persistence of cancer as a leading public health challenge reflects its profound biological complexity, and the coordinated interplay of hallmarks, including uncontrolled proliferation, immune evasion, resistance to cell death, and metabolic reprogramming. ^[1,2]^ Among these hallmarks, metabolic reprogramming is distinctive in that it not only sustains the energetic and biosynthetic demands of proliferating cells ^[3,4]^ but also modulates other hallmarks—particularly immune evasion and resistance to regulated cell death (RCD) mechanisms triggered by metabolic stress. ^[5–7]^

Metabolic reprogramming plays a central role in immunosuppression, either by limiting essential nutrients to immune cells or through the accumulation of immunosuppressive metabolites such as lactate and kynurenine. ^[6,8]^ Metabolic plasticity enables cancer cells to resist specific RCD pathways, such as apoptosis and ferroptosis, and more recently described metabolic forms of cell death, such as cuproptosis and disulfidptosis, through adaptations including the Warburg effect, glutamine dependence, or lipid remodeling. ^[5,9]^ These adaptations generate metabolic dependencies that constitute therapeutic vulnerabilities exploitable through pathway-specific inhibition. ^[10]^

The metabolic alterations underlying metabolic reprogramming arise from (epi)genomic and transcriptomic modifications in genes encoding enzymes and their non-coding RNA regulators across multiple metabolic pathways.^[11–13]^ However, despite the availability of extensive multi-omic datasets, systematic integrative analyses linking metabolic genes, non-coding regulators, tumor phenotypes, prognosis, and tumor immune context remain limited. ^[14,15]^

The regulatory logic that coordinates metabolic pathways with non-coding RNA control, tumor phenotypes, immune ecologies, and clinical outcomes remains largely unresolved, and no existing framework systematically reconstructs these relationships across multi-omic layers and cancer types.

Building on our previous work on the multi-omic construction of RCD signatures, ^[16]^ we now extend this strategy to the metabolic domain by integrating enzyme-coding genes and their non-coding regulators across 127 Kyoto Encyclopedia of Genes and Genomes (KEGG) pathways and seven metabolic categories. ^[17]^ This analysis encompasses 33 cancer types from The Cancer Genome Atlas (TCGA). ^[18]^ This foundation enables the systematic derivation of multi-omic, phenotypic, and clinical relationships that underpin metabolic reprogramming at scale.

In this study, we introduce the concept of omic-specific metabolic signatures, defined as multidimensional units that integrate metabolic-pathway context with phenotypic, prognostic, and immune attributes. These signatures capture coordinated cross-layer behavior rather than simple co-expression, embedding molecular and clinical information into structured metabolic states. This formulation provides the foundation required to identify how metabolic programs are regulated by shared upstream non-coding RNAs and how these regulatory influences align or oppose one another across tumor contexts.

Using this integrative foundation, we developed OncoMetabolismGPS, a multi-omic analytical framework that identifies omic-specific metabolic signatures, maps their shared upstream regulatory interactions, and classifies each regulator–signature pair as convergent or divergent across molecular, phenotypic, immune, and clinical dimensions. This strategy reconstructs a Pan-Cancer atlas of metabolic regulatory circuitries, enabling the systematic identification of aligned versus opposing regulatory influences within metabolic pathways. The full workflow, including data layers, signature construction, regulatory interrogation, and circuitry classification, is summarized in Figure 1.

**Figure 1.**
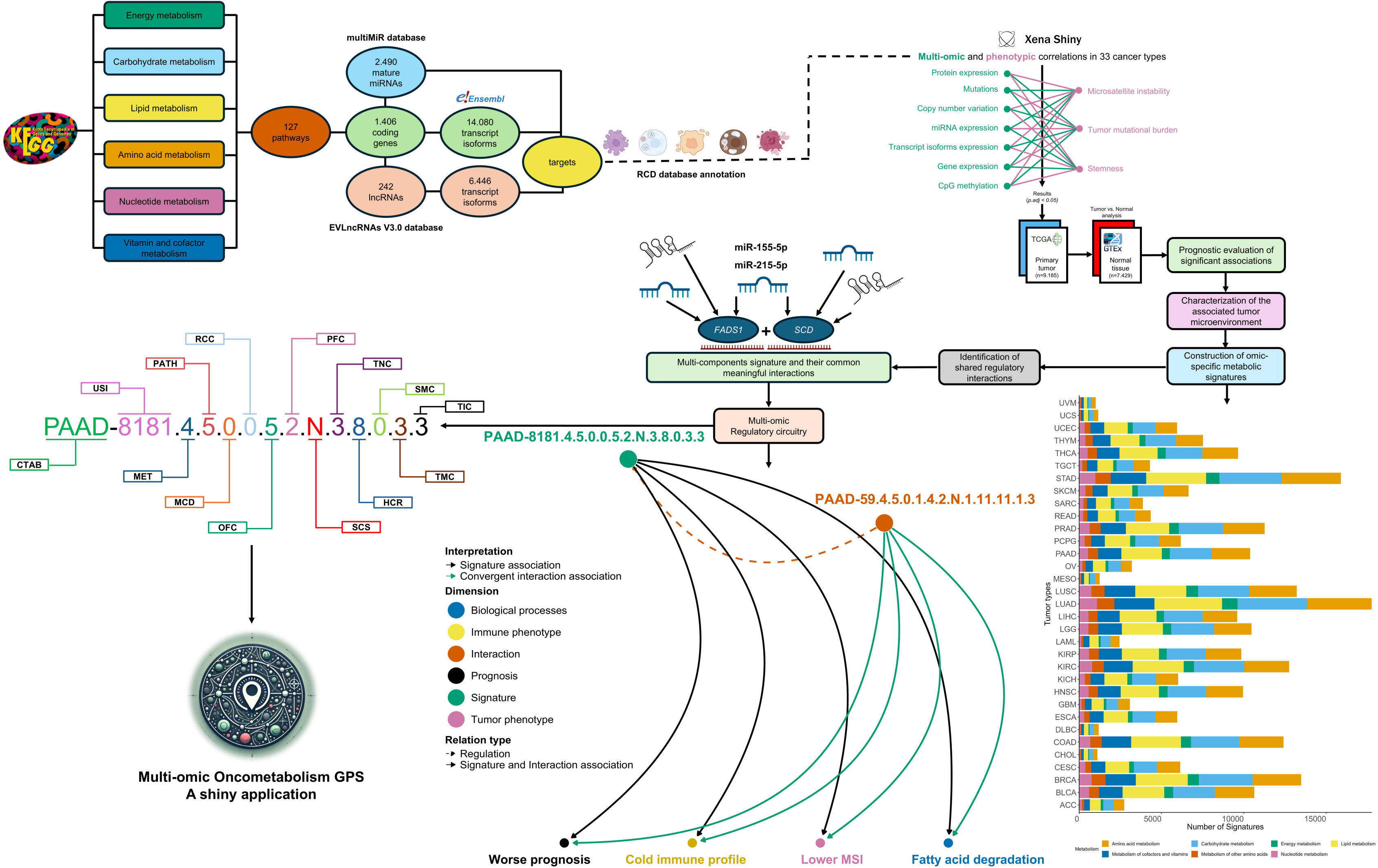
Overview of the OncoMetabolismGPS framework for the identification, integration, and interpretation of multi-omic metabolic signatures in cancer. The workflow begins with the selection of enzyme-coding genes involved in KEGG-defined metabolic pathways, together with their experimentally validated regulatory interactions with miRNAs and lncRNAs (miRTarBase and EVlncRNAs 3.0). These molecular components are annotated for participation in regulated cell death (RCD) mechanisms. Next, multi-omic, phenotypic, and clinical data from 33 TCGA tumor types are integrated. Significant associations are then evaluated for their clinical impact, and immune microenvironment features inferred by immune-cell deconvolution. Based on these integrative associations, omic-specific metabolic signatures are constructed within each cancer type, defined by shared metabolic function, phenotypic directionality, prognostic coherence, and immune-context compatibility. For each signature, shared upstream regulators are identified, generating integrated regulatory circuitries that are subsequently classified as convergent or divergent according to their cross-dimensional behavior. Each signature is assigned a standardized 13-token nomenclature encoding tumor type, omic layer, metabolic pathway, RCD involvement, phenotypic class, prognostic direction, and immune microenvironment profile. Collectively, these analytical components make up the OncoMetabolismGPS framework, which functions as a multi-omic “Guided Positioning System” for tumor metabolism. It systematically positions every metabolic signature within a multidimensional molecular-to-clinical landscape, enabling the identification of regulatory circuitries that link metabolic pathways to phenotypic and immune outcomes. All resulting signatures, interactions, and evaluation metrics are accessible through the interactive OncoMetabolismGPS Shiny application described in Section 2.10.

Finally, we establish a standardized nomenclature system, and we implement all findings in the OncoMetabolismGPS Shiny application (https://OncoMetabolismGPS.shinyapps.io/Multi-omicOncoMetabolismGPSShiny/). This interactive multi-omic atlas enables users to visualize metabolic signatures, explore regulatory circuitries, and construct custom signature sets according to molecular, phenotypic, or immunological criteria.

Together, these components establish the first unified framework for reconstructing metabolic regulatory circuitries across cancers, enabling systematic exploration of how metabolic pathways interface with non-coding RNA regulation, tumor phenotypes, immune ecologies, and clinical outcomes.

## 2. Results

### 2.1. Molecular targets identified in metabolic processes and annotation in metabolic cell death mechanisms

We retrieved 1,406 unique enzyme-coding genes functionally associated with catalytic reactions distributed across 127 KEGG-defined metabolic pathways encompassing anabolic and catabolic processes. These pathways were grouped into seven major metabolic categories: carbohydrate metabolism, lipid metabolism, energy metabolism, amino-acid metabolism, metabolism of other amino acids, nucleotide metabolism, and metabolism of cofactors and vitamins (Table S1). For these enzyme-coding genes, we identified 2,490 microRNAs (miRNAs) and 242 long non-coding RNAs (lncRNAs) with experimentally validated regulatory interactions (Tables S2–S3). For each enzyme or regulator, 20,526 transcript isoform identifiers (GRCh38.p13) were retrieved from Ensembl (Tables S4–S5).

Among these targets, 309 molecules were annotated to regulated metabolic cell death mechanisms, including alkaliptosis, apoptosis, autophagy-dependent cell death, cuproptosis, entotic cell death, ferroptosis, lysosome-dependent cell death, necroptosis, oxeiptosis, parthanatos, and pyroptosis (Table S6).

### 2.2. Multi-omic and phenotypic associations across 33 cancers

We developed an R-based computational pipeline to perform approximately 2.94 million association tests across multiple omic layers—bulk-RNA gene expression, transcript expression, miRNA expression, CpG methylation, copy number variation (CNV), somatic mutation, and protein expression—evaluating their relationships with three phenotypic variables: microsatellite instability (MSI), tumor mutational burden (TMB), and tumor stemness (TSM). Analyses were conducted for molecular targets comprising enzyme-coding genes, miRNAs, lncRNAs, and transcript isoforms across 33 TCGA cancer types.

Overall, we identified 171,782 statistically significant associations (adjusted p < 0.05) across all omic layers, phenotypic variables, and cancer types (Dataset S1). Transcript-isoform expression accounted for 51.7% of these associations, followed by gene expression (15.4%), CpG methylation (14.7%), somatic mutation (7.5%), miRNA expression (6.3%), and CNV (4.4%). Significantly fewer correlations were observed for protein expression. By phenotypic variable, TSM contributed 72.1% of associations, TMB 14.3%, and MSI 13.6%.

After annotation to metabolic pathways, metabolic cell death mechanisms, and regulatory interactions, the number of associations expanded—through legitimate multi-mapping—to 463,433 entries (Dataset S2), reflecting the intrinsic multifunctionality of metabolic genes and their non-coding RNA regulators.

### 2.3. Prognostic relevance and tumor microenvironment profiles of significant associations

Of the 463,433 associations, 183,136 (39.5%) exhibited prognostic relevance in Cox regression or Kaplan–Meier analysis. Among these prognostic associations, TME classes were distributed as dual 50.7%, anti-tumoral 23.9%, pro-tumoral 17.2%, and non-significant 8.2%. Immune-phenotype classification revealed a predominance of cold profiles (81.9%), followed by hot 12.4% and variable 5.7% (Dataset S2).

### 2.4. Construction of omics-specific metabolic signatures

To integrate multi-omic and phenotypic associations into biologically interpretable entities, we constructed omic-specific metabolic signatures within each cancer type. Each signature is mono-omic, comprising one or more elements from the same omic layer—enzyme-coding genes, miRNAs, lncRNAs, or transcript isoforms—that (i) share a common KEGG-defined metabolic pathway and the same metabolic cell death mechanism; (ii) exhibit concordant associations with tumor phenotypes and prognostic outcomes across all survival endpoints; and (iii) align with the same immune microenvironment class.

Elements displaying fully concordant profiles across molecular, phenotypic, prognostic, and immune dimensions were grouped into composite signatures, whereas unmatched gene attributes were kept as single-member signatures. Because a good deal of enzyme-coding genes take part in multiple pathways or cell death mechanisms, individual gene attributes may recur in distinct context-specific signatures. We identified 241,415 signatures across 33 tumor types (Dataset S3).

By omic layer, signatures were distributed: miRNA expression, 102,272 (42.4%); transcript-isoform expression, 65,590 (27.2%); gene expression, 31,694 (13.1%); CpG methylation, 25,400 (10.5%); CNV, 8,608 (3.6%); somatic mutation, 7,827 (3.2%); and protein expression, 24 (<0.1%) (Figure 2A).

**Figure 2.**
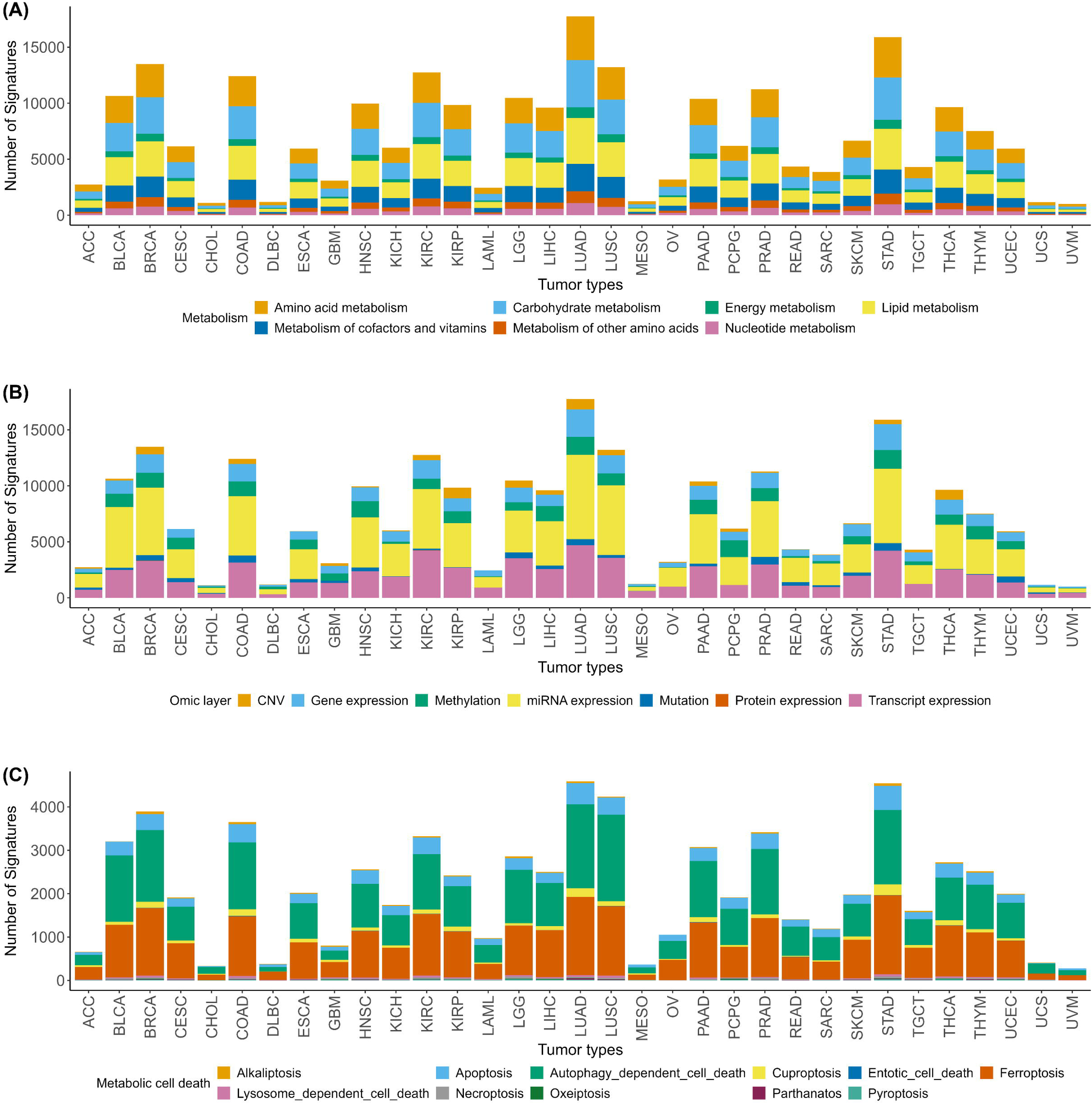
Distribution of omic-specific metabolic signatures across tumor types, molecular layers, and metabolic contexts. (A) Distribution of signatures by metabolic category across 33 TCGA tumor types, including amino acid metabolism (56,811), lipid metabolism (60,070), energy metabolism (12,721), carbohydrate metabolism (60,860), metabolism of cofactors and vitamins (35,248), metabolism of other amino acids (15,064), and nucleotide metabolism (14,774). (B) Distribution by omic layer: gene expression (31,694), CpG methylation (25,400), miRNA expression (102,272), protein expression (24), mutation (7,827), CNV (8.608), and transcript expression (79,723). (C) Distribution by metabolic cell death mechanism (total > 70,000 signatures), including autophagy-dependent cell death (29,701), ferroptosis (30,021), apoptosis (7,842), lysosome-dependent cell death (772), cuproptosis (2,602), alkaliptosis (738), necroptosis (645), pyroptosis (403), oxeiptosis (140), parthanatos (204), and entotic cell death (101).

By metabolic category, signatures were distributed: carbohydrate metabolism, 57,563 (23.8%); lipid metabolism, 56,870 (23.5%); amino-acid metabolism, 53,422 (22.1%); metabolism of cofactors and vitamins, 33,485 (13.8%); metabolism of other amino acids, 14,186 (5.8%); nucleotide metabolism, 13,831 (5.7%); and energy metabolism, 12,058 (5.0%) (Figure 2B).

Over 70,000 signatures were mapped to metabolic cell death mechanisms, primarily autophagy-dependent cell death, 29,142 (41.3%), and ferroptosis, 28,529 (40.4%), followed by apoptosis, 7,626 (10.8%); the remaining mechanisms were less frequent (Figure 2C).

### 2.5. Signatures association with tumor phenotype, clinical outcome, and tumor microenvironment profile

Having established the omic-specific metabolic signatures, we next examined whether they share upstream regulatory molecules that coordinate metabolic behavior. Signatures showed significant associations with MSI, TMB, and, most prominently, TSM across all omic layers, metabolic categories, and metabolic cell death mechanisms (Figure S1; Tables S9–S11). Most associations corresponded to dual phenotypic profiles, followed by pro-tumoral and anti-tumoral signatures (Figure S2; Tables S12–S14). When examining immune phenotypes, cold-associated signatures predominated across datasets, whereas hot and variable immune profiles represented smaller fractions (Figure S3; Tables S15–S17).

A comprehensive summary of counts and percentages for each omic layer, metabolic category, and metabolic cell death mechanism is provided in Tables S9–S17, while the complete dataset of individual associations for all signatures and tumor types is available in Dataset S3.

Regarding the clinical endpoints, the prognostic associations—capturing both risk-enhancing and protective effects—are presented in Tables S18–S29 and summarized in Figures S4–S7.

### 2.6 Signature nomenclature

To enable unambiguous identification and cross-referencing of all signatures, we implemented a structured nomenclature system adapted from our previously published framework for RCD–RCD-associated signatures ^[16]^ and extended for the metabolic context of the present study. The original RCD system contained eleven hierarchical tokens encoding tumor, omic, phenotypic, prognostic, and immune attributes. In the OncoMetabolismGPS framework, this structure was expanded to a 13-component code that now also captures the metabolic and regulatory dimensions specific to this framework (Figure 3; Supporting Information -Method C).

**Figure 3.**
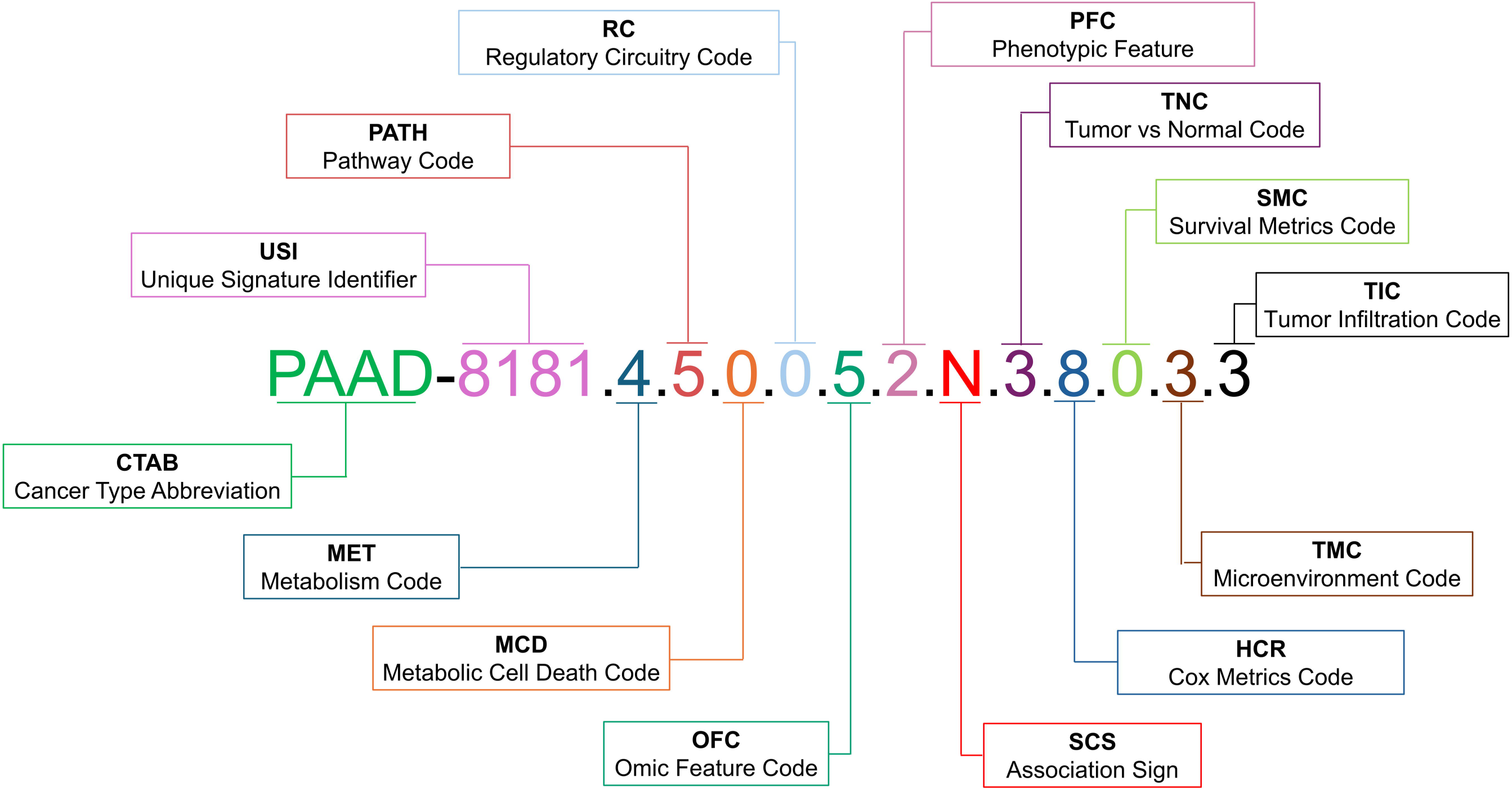
Hierarchical nomenclature system of omic-specific metabolic signatures in OncoMetabolismGPS. This figure illustrates the extended nomenclature and coding structure used to uniquely identify and rank multi-omic metabolic signatures. Each signature is encoded by a 13-token alphanumeric string that integrates tumor context, metabolic and regulatory dimensions, phenotypic behavior, and immune classification. The coding sequence follows the format: **[CTAB] [USI].[MET].[PATH].[MCD].[RCC].[OFC].[PFC].[SCS].[TNC].[HRC]. [SMC].[TMC].[TIC**]. Here, CTAB is the TCGA tumor abbreviation (cancer abbreviations defined in Table S7), USI the unique series identifier, MET the metabolic category (e.g., lipid, amino acid, carbohydrate), PATH the pathway index within that category, MCD the metabolic cell death flag, INT the shared-interaction flag showing common regulators, OFC and PFC the omic and phenotypic layer codes, SCS the correlation sign (positive or negative), TNC the tumor-versus-normal expression code, HRC and SMC the hazard-direction and survival worst-group templates across the survival metrics DSS (disease-specific survival), DFI (disease-free interval), PFI (progression-free interval), and OS (overall survival), TMC the tumor-microenvironment class (anti-tumoral, dual, or pro-tumoral), and TIC the immune phenotype (hot, cold, or variable). Relative to the earlier nomenclature used for RCD-associated signatures [16], three new fields are introduced: MET (metabolism code), PATH (pathway code), and RCC (regulatory circuitry code), extending the system from 11 to 13 components.

Each signature identifier follows the general token format

**[CTAB]-[USI].[MET].[PATH].[MCD].[RCC].[OFC].[PFC].[SCS].[TNC].[HRC]. [SMC].[TMC].[TIC]** and encodes, in compact form, its tumor type, metabolic category, omic layer, phenotype association, survival behavior, and immune classification (Supporting Information -Method C). Three new fields were introduced relative to the prior nomenclature:

**[MET]** (Metabolism code) shows the metabolic domain (e.g., lipid, amino acid, carbohydrate).

**[PATH]** (Pathway code) denotes the KEGG pathway index within each metabolic domain.

**[RCC]** (Regulatory Circuitry code) identifies whether the signature takes part in a convergent or divergent regulatory pair within its tumor context.

Together, these tokens provide a lossless, human- and machine-readable representation of each signature’s biological context and analytical metadata. Figure 6 illustrates the hierarchical logic and an example of a fully expanded identifier, while complete token definitions and codebooks are available in Supporting Information -Method C.

Because the signature compendium (Dataset S3) includes both single- and multi-element configurations, and because individual metabolic components may take part in different biological contexts, the same molecular elements can appear in distinct signatures across tumor types whenever they satisfy layer-specific, phenotypic, prognostic, and immune-context criteria. The *AKR1C1* + *AKR1C2* pair illustrates this principle: these genes formed tumor-specific signatures in LUAD, GBM, KIRP, PAAD, PRAD, SKCM, THYM, UCS, and KIRC (abbreviations in Table S7), each reflecting coherent associations at both the gene-expression and DNA-methylation layers. In prostate cancer, for example, the PRAD-specific signature (PRAD-1119.4.13.6.0.6.3.N.2.5.0.1.3) consistently mapped to a cold immune microenvironment and exhibited an adverse prognostic profile, with significant effects in overall survival (p = 0.0136) and disease-specific survival (p = 0.00318). This case shows how individual metabolic genes can give rise to distinct, cancer-type-specific metabolic signatures, each shaped by its surrounding phenotypic, clinical, and immune context.

### 2.7. Shared molecular regulatory interactions within signatures

This step serves as a bridge between metabolic signatures and their regulatory circuitry. We next examined whether the components of each signature were connected through shared molecular regulators. For enzyme-coding gene signatures, we intersected experimentally validated miRNA–mRNA and lncRNA–mRNA interactions, keeping only those regulators common to all members. For miRNA- and lncRNA-based signatures, we applied the equivalent criterion in the reverse regulatory direction, identifying coding genes that were co-targeted by all non-coding members.

Of the 241,415 total signatures, 57,033 (23.6%) were multi-member, whereas the remaining 184,382 (76.4%) comprised single elements. Among the multi-member signatures, 11,021 (4.6%) were linked by at least one shared regulator, whereas 46,012 (19.1%) showed no common regulatory connection among their members. Among the single-member signatures, 171,084 (70.9%) were connected to at least one common regulatory molecule, while 13,298 (5.5%) had no known regulatory connection. In total, 182,105 signatures (75.4%) exhibited at least one molecular regulatory interaction across cancer types.

As an illustrative example, in PAAD, the metabolic signature PAAD-1340.5.4.0.1.6.3.N.3.3.0.3.3, composed of *NNT* and *SIRT4*, both involved in nicotinate and nicotinamide metabolism within the cofactors-and-vitamins metabolic category, shared five experimentally validated upstream regulators—hsa-miR-103a-3p, hsa-miR-15a-5p, hsa-miR-15b-5p, hsa-miR-16-5p, and hsa-miR-195-5p. The presence of this multi-miRNA set illustrates how multi-element metabolic signatures can be connected through multiple common regulatory molecules, highlighting the integrative nature of regulatory interactions within PAAD.

### 2.8. Meaningful convergent and divergent regulatory relationships between shared regulators and signatures

This section makes up the core output of the framework: a multi-omic atlas of convergent and divergent metabolic regulatory circuitries reconstructed across 33 cancer types. To characterize the directionality of regulatory influence, we classified each regulator–signature pair as either convergent or divergent based on the sign of their association across molecular, phenotypic, and clinical dimensions. A convergent relationship occurs when the regulator and its target signature exhibit correlated behavior, acting in the same biological direction across phenotypic and prognostic contexts. Conversely, a divergent relationship reflects opposite association directions, implying regulatory compensation or feedback within the metabolic circuitry.

For signatures with shared regulators, we compared the direction of association between each signature and its regulator across tumor phenotypes (MSI, TMB, TSM), survival metric outcomes (overall survival (OS), disease-specific survival (DSS), disease-free interval (DFI), and progression-free interval (PFI)), and immune-infiltration profiles (hot, cold, variable).

This analysis identified a large subset of signatures engaged in measurable regulatory interactions across these dimensions. In total, 84,270 signatures exhibited at least one molecular regulatory interaction. Because a single signature can be linked to multiple regulators, and individual regulators can interact with several signatures, the total number of regulator–signature relationships analyzed for convergence and divergence exceeded the number of unique signatures, totaling 204,591 distinct interactions (Dataset S4).

Across all regulatory interactions, 27.2% (55,759) displayed only divergent associations, while 24.4% (49,856) exhibited only convergent associations. Among domain-specific analyses, interactions involving the immune profile accounted for the largest proportion (48.4%, 98,976), followed by those associated with tumor phenotypes (34.2%, 70,072), Cox regression endpoints (29.3%, 59,980), and overall survival outcomes (21%, 43,046). These results show that divergent regulatory behavior predominates, implying that shared regulators often exert context-dependent effects across tumor phenotypes, immune profiles, and clinical outcomes.

In LGG, a lipid-metabolism signature composed of *FADS1* and *SCD*, annotated to the biosynthesis of unsaturated fatty acids and the ferroptosis pathway, exhibited a divergent regulatory pattern. The metabolic signature LGG-1494.4.2.6.1.6.3.P.3.8.8.2.3 showed a positive association with TSM through its gene-expression layer (ρ = 0.216), corresponded to a cold immune microenvironment, and was linked to a protective prognostic effect in Cox analyses. In contrast, its shared upstream regulators—hsa-miR-215-5p and hsa-miR-155-5p, displayed opposite trends, including negative correlations with TSM (ρ = –0.529 and -0.253, respectively), associations with a hot immune phenotype, and a risky prognostic profile (Figure 4A; Dataset S4). Together, the LGG-1494.4.2.6.1.6.3.P.3.8.8.2.3 and its shared upstream regulators exemplifies a divergent metabolic regulatory circuitry, in which the regulatory layer acts in an opposing biological direction relative to the metabolic signature.

**Figure 4.**
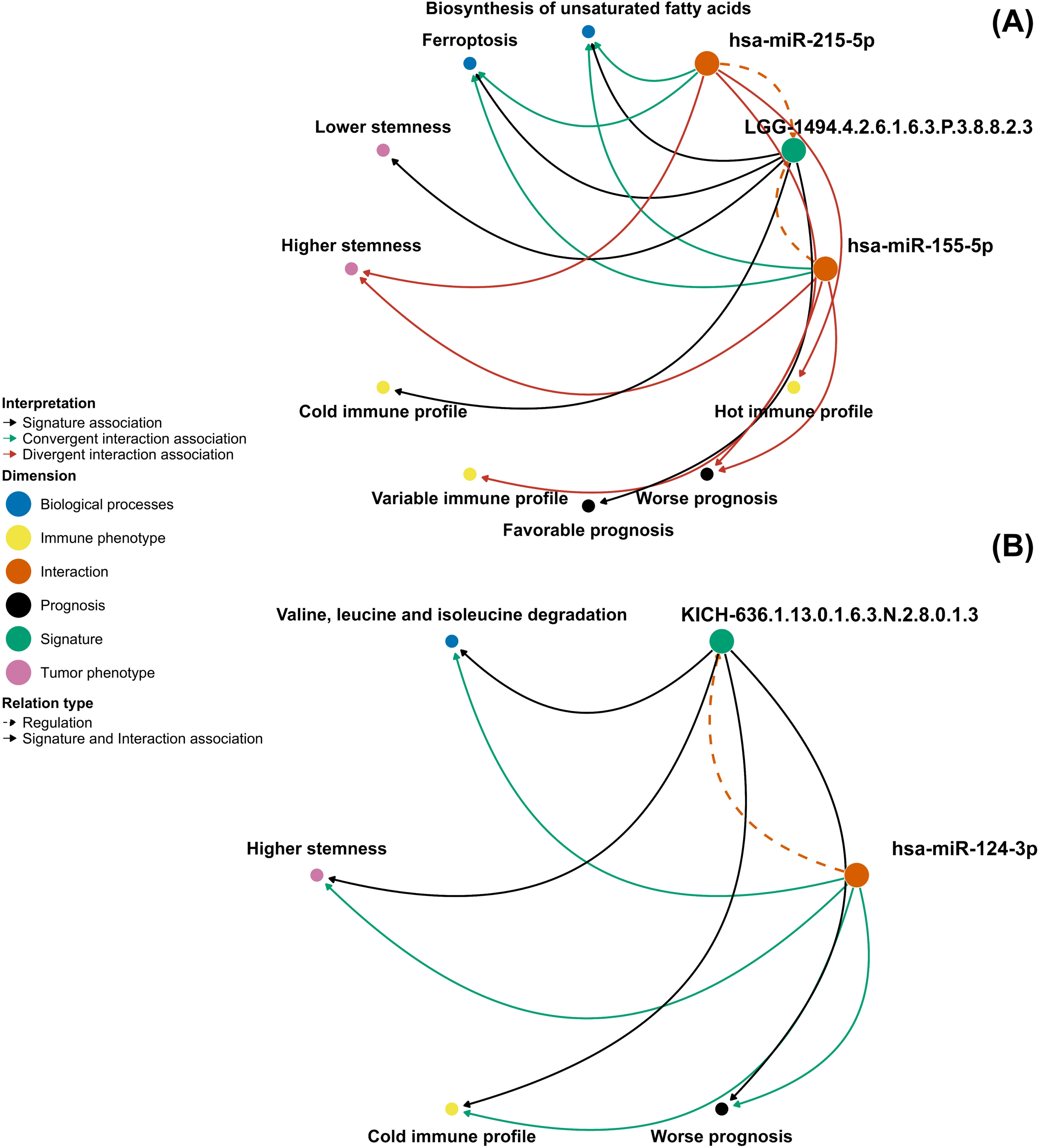
Convergent and divergent regulatory circuitries across tumor contexts. (A) In LGG, a lipid-metabolism signature composed of *FADS1* and *SCD*—annotated to the biosynthesis of unsaturated fatty acids and ferroptosis—formed the metabolic signature LGG-1494.4.2.6.1.6.3.P.3.8.8.2.3, which displayed a divergent regulatory circuitry relative to its paired regulatory interactions (comprising miR-215-5p and miR-155-5p). The metabolic signature was associated with higher TSM, a cold immune microenvironment, and a favorable prognosis, whereas the regulatory signature showed the opposite pattern—lower TSM, hot immune profiles, and poor prognosis. This antagonistic configuration illustrates how divergent circuitries encode metabolic–immune trade-offs, in which metabolic advantage and immune visibility are regulated in opposing directions within the same tumor context. (B) In KICH, a metabolic signature composed of *ACAT2* and *HADH*, linked to branched-chain amino-acid degradation, tryptophan metabolism, and the butanoate pathway, formed the metabolic signature KICH-636.1.13.0.1.6.3.N.2.8.0.1.3 and exhibited convergent regulation with its paired regulatory interaction (defined by miR-124-3p). Both signatures showed negative correlations with TSM, association with cold immune profiles, and unfavorable prognostic behavior, demonstrating alignment across molecular, phenotypic, and clinical dimensions. Together, panels (A) and (B) illustrate how divergent circuitries mark axes of adaptive metabolic–immune plasticity, whereas convergent circuitries stabilize coordinated metabolic programs within tumor contexts.

In KICH, a metabolic signature composed of *ACAT2* and *HADH*—annotated to branched-chain amino acid degradation pathway—exhibited a convergent regulatory pattern. The metabolic signature KICH-636.1.13.0.1.6.3.N.2.8.0.1.3 showed a negative association with TSM (ρ = –0.52), corresponded to a cold immune microenvironment, and was linked to a risky prognostic effect in Cox analyses. Its shared regulator, hsa-miR-124-3p displayed the same directional behavior, including a negative correlation with TSM (ρ = –0.45), a cold immune phenotype, and an adverse prognostic profile. This coherence across molecular, phenotypic, immune, and clinical dimensions defines a convergent metabolic regulatory circuitry, in which both the metabolic and regulatory signatures act in the same biological direction within the tumor context (Figure 4B; Dataset S4).

### 2.9. Multi-omic metabolic regulatory circuitries

In our analytical framework, a metabolic regulatory circuitry is operationally defined as a cancer type-specific pair of omic signatures in which one signature comprises enzyme-coding genes involved in metabolic pathways and the other comprises non-coding RNA regulators (miRNAs or lncRNAs) with experimentally validated interactions with those genes. For each pair, all components of the regulatory signature are linked to all components of the target signature through validated molecular interactions, ensuring complete cross-connectivity. These paired signatures were analyzed for directionality of association across molecular, phenotypic, and clinical dimensions to determine whether their regulatory relationship was convergent, when both signatures exhibited aligned associations with tumor features (e.g., both linked to risk or both to protection), or divergent, when their associations were inverse (e.g., one enriched in protective phenotypes while the other corresponded to risk). Circuitries identified in this manner represent structured regulatory modules coupling metabolic gene programs to their non-coding regulators within defined tumor contexts.

From the 204,591 distinct regulator–signature interactions identified in Dataset S4, we generated composite interaction signatures and compared them against the catalog of omic-specific metabolic signatures defined previously (Dataset S3) to determine overlaps. This mapping revealed 24,796 matching pairs between the two datasets (Dataset S5).

Exploratory analysis of the signature–interaction pairs revealed that the vast majority (23,288 pairs; ∼96.9%) were classified as unique in both datasets, showing that each signature and its corresponding interaction involved a single, specific molecular element. A smaller subset (904 pairs; ∼3.8%) comprised unique signatures linked to multiple interactions, while 568 pairs (∼2.4%) exhibited the opposite pattern—multiple molecules within the signature but a single molecule in the interaction. Only 36 pairs (<0.2%) contained multiple molecules in both the signature and the interaction, representing multifactorial regulatory associations.

To illustrate a representative example, we selected a pair containing multiple molecules in both the signature and the interaction, representing convergent associations (Figure 5). The signature PAAD-8181.4.5.0.0.5.2.N.3.8.0.3.3, specific to the fatty acid degradation pathway, is composed of the transcripts ENST00000503281 and ENST00000553117 of the *ALDH7A1* (Aldehyde Dehydrogenase 7 Family Member A1) gene and functionally integrates with the miRNA interaction hsa-let-7b-5p + hsa-miR-193b-3p (PAAD-59.4.5.0.1.4.2.N.1.11.11.1.3). In PAAD patients, this association is characterized by a convergent relationship between transcript and miRNA expression layers, both correlated with the MSI phenotype. The negative correlations observed for the signature (ρ = −0.46; padj= 0.0064) and for the interaction (ρ = −0.38; padj= 5.3×10⁻⁶) show a coordinated regulatory effect. In survival analyses, both the signature and the interaction exhibited a high-risk profile in a convergent manner across DSS, DFI, and OS survival metrics, demonstrating prognostic coherence between the omic layers. For the transcriptomic signature, p-values were 0.0264 (OS), 0.0293 (DSS), and 0.0198 (DFI), showing significant associations with worse survival outcomes. For the miRNA interaction, the associations were even stronger, with p = 8.13×10⁻⁵ (OS), p = 5.15×10⁻⁴ (DSS), and p = 4.07×10⁻⁴ (DFI). From an immunological and microenvironmental perspective, both layers displayed concordant profiles, being associated with a pro-tumoral and immunologically cold microenvironment, consistent with reduced immune infiltration and unfavorable clinical outcome.

**Figure 5.**
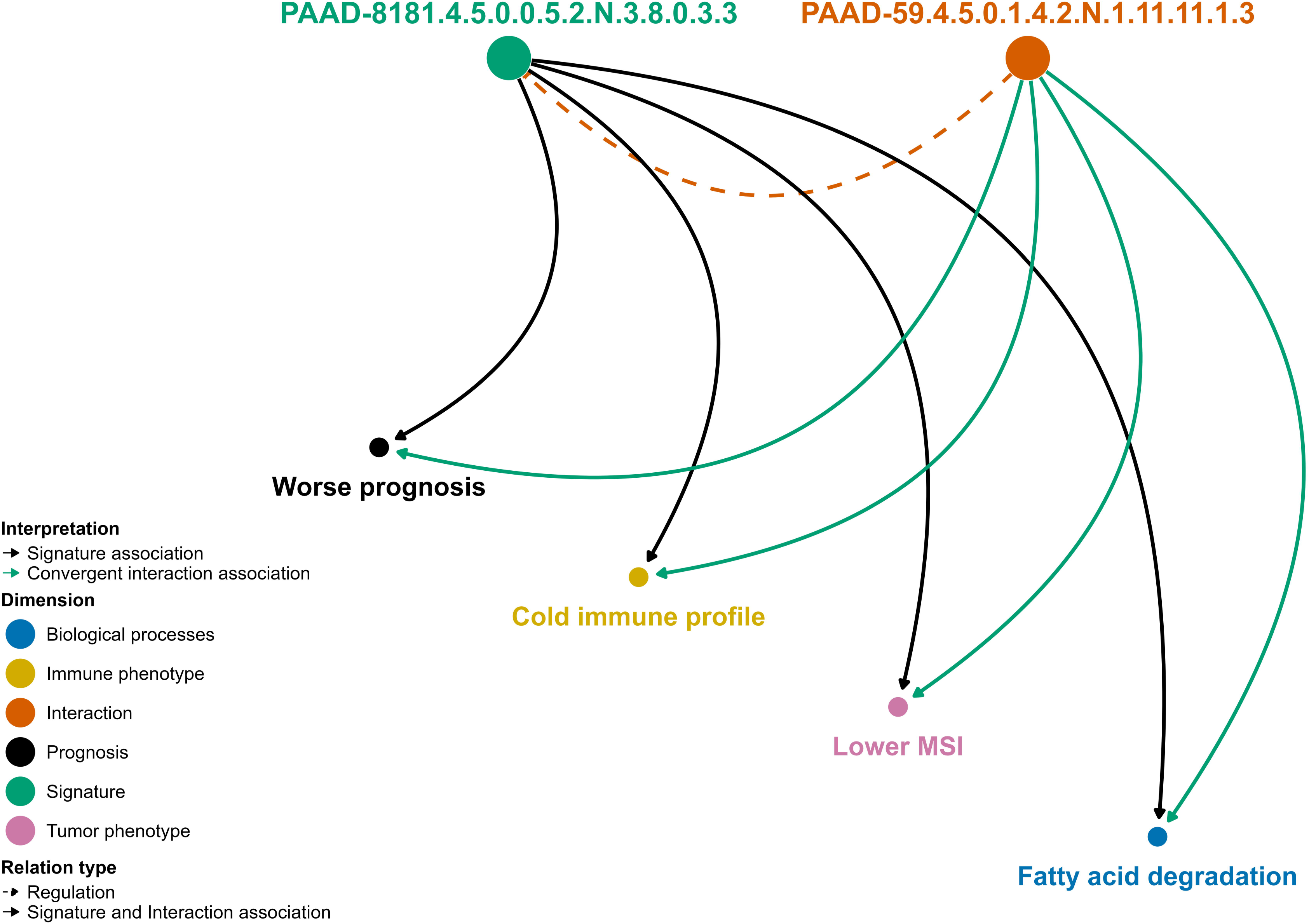
Convergent metabolic regulatory circuitry in PAAD. In PAAD, a multi-element transcriptomic signature PAAD-8181.4.5.0.0.5.2.N.3.8.0.3.3—composed of *ALDH7A1* transcripts ENST00000503281 and ENST00000553117 and annotated to the fatty-acid degradation pathway—forms a convergent circuitry with the paired regulatory signature PAAD-59.4.5.0.1.4.2.N.1.11.11.1.3, comprising hsa-let-7b-5p and hsa-miR-193b-3p. Both signatures showed negative associations with MSI (ρ = –0.46 and –0.38) and displayed coherent high-risk prognostic profiles across OS, DSS, and DFI survival metrics. Immunologically, each signature corresponded to a pro-tumoral, cold microenvironment, indicating reduced immune infiltration. This alignment across molecular, phenotypic, survival, and immune dimensions exemplifies a convergent metabolic regulatory circuitry in which metabolic and regulatory signatures act in the same biological direction within the tumor context.

### 2.10. OncoMetabolismGPS Shiny application for interactive exploration

To facilitate access, navigation, and interpretation of the multidimensional metabolic signatures generated in this study, we developed OncoMetabolismGPS, an interactive Shiny-based web application (https://OncoMetabolismGPS.shinyapps.io/Multi-omicOncoMetabolismGPSShiny/). The acronym OncoMetabolismGPS encapsulates both the structural logic and navigational philosophy of the framework. Analogous to a Guided Positioning System, the platform situates each omic-specific metabolic signature within a multidimensional coordinate space defined by molecular, phenotypic, immune, and clinical attributes. The structured 13-token nomenclature functions as a geospatial code or zip-code-like identifier that encodes the biological location of each signature within this landscape, enabling precise positioning and contextual comparison across tumor types. Rather than a static catalog of associations, the framework operates as a metabolic geolocation atlas in which signatures, pathways, and regulatory circuitries can be dynamically explored and cross-referenced.

The OncoMetabolismGPS Shiny application operationalizes this principle by providing an interactive interface that allows users to navigate the metabolic terrain of cancer. Through layered visualization modules, users can traverse from single enzyme-coding genes or non-coding regulators to higher-order metabolic signatures and their convergent or divergent regulatory circuitries, examining how molecular patterns align with phenotypic, prognostic, and immune dimensions across tumor contexts. The interface integrates all omic-specific metabolic signatures with their associated metabolic pathways, RCD annotations, tumor phenotypes (MSI, TMB, TSM), survival outcomes (OS, DSS, DFI, PFI), immune infiltration profiles, and shared regulatory circuitries.

Users can query signatures by cancer type, omic layer, metabolic pathway, cell death mechanism, phenotypic association, immune microenvironment class, or specific gene, miRNA, or lncRNA identifiers. For each selected signature, the interface displays its molecular components, the regulatory interactions that define convergent or divergent circuitries, association statistics across phenotypic and clinical dimensions, immune microenvironment classification, and downloadable tables and visual summaries. The application provides an operational framework for exploring how metabolic reprogramming shapes tumor behavior across biological contexts and serves as a resource for hypothesis generation, pathway prioritization, identification of metabolic vulnerabilities, and selection of biomarker candidates for translational validation (Public access and source code are available in the associated GitHub repository.)

## 3. Discussion

### 3.1. Integrated View of Metabolic Reprogramming and Cell-Fate Control

Metabolic rewiring, immune modulation, and RCD constitute interdependent axes that collectively dictate tumor behavior. ^[4,5,9]^ Metabolic programs intersect with apoptotic, ferroptotic, autophagic, and necrotic pathways, influencing cell-fate decisions in a context-dependent manner. These same metabolic alterations promote immune evasion through nutrient competition, redox remodeling, and disrupted antigen presentation, while microenvironmental stressors—hypoxia, ROS accumulation, and amino-acid limitation—activate specific RCD modules that feed back into tumor evolution.

The metabolic–immune–RCD axis is further shaped by local environmental constraints such as pH, nutrient gradients, and stromal interactions, as illustrated in injury models where metabolic stress elicits distinct RCD patterns depending on substrate availability. ^[4,5,9]^ Within this integrated landscape, tumor metabolism functions as a central regulatory hub that links survival, death, and immune escape programs. This systems-level view motivated the development of our multi-omic framework designed to reconstruct metabolic signatures and their governing regulatory logic across diverse cancers.

The integration between cellular metabolism and RCD mechanisms shows that metabolic pathways do not act in isolation. They form a multifunctional system in which metabolic enzymes take part in various physiological and adaptive processes, conferring plasticity to cellular responses. ^[19]^ The regulation of these mechanisms depends strongly on the metabolic state. Processes such as ferroptosis and autophagy are sensitive to the balance between nutrient availability, oxidative stress, and the efficiency of bioenergetic routes. For example, in cases of acute kidney injury, the energetic metabolism of tubular epithelial cells is reprogrammed, involving coordinated changes in lipid, glucose, and amino acid metabolism. This reprogramming is associated with the activation of different forms of cell death—autophagy, apoptosis, necroptosis, pyroptosis, and ferroptosis—establishing a metabolism–cell death axis that influences both tissue progression and repair. ^[20]^ Similarly, in colorectal carcinomas, oncogene-driven metabolic reprogramming alters glycolysis, glutaminolysis, one-carbon metabolism, and lipid pathways. These changes support cell proliferation, immune evasion, and therapeutic resistance, while the tumor microenvironment both modulates and is modulated by these routes. ^[21]^

Alternative forms of metabolic cell death, such as cuproptosis, reinforce this interdependence: therapies that redirect tumor metabolism from a glycolytic to a more oxidative state increase sensitivity to copper-induced toxicity, intensifying cell death. ^[22]^

Taken together, these observations indicate that the metabolic state determines cellular vulnerabilities and fate. Based on this, this study proposes a multidimensional Pan-Cancer framework that integrates metabolic, phenotypic, and clinical information to identify signatures and molecular interactions within a regulatory circuitry. This framework, implemented in the OncoMetabolismGPS application, demonstrates that tumor metabolism has a layered regulatory architecture in which biochemical activity, RCD mechanisms, tumor aggressiveness, and immune response are interconnected. This approach enables the identification of context-dependent metabolic vulnerabilities that can serve as strategic points for future basic and applied investigations with therapeutic relevance.

This study establishes a multidimensional framework that integrates metabolic, phenotypic, and clinical information to explain how metabolic reprogramming organizes tumor behavior across cancers.

### 3.2. A Pan-Cancer Landscape of Multi-Omic Metabolic Signatures

Across 33 tumor types, we constructed a unified atlas of metabolic signatures that integrates multi-omic, phenotypic, immunological, and prognostic information. These signatures define coherent metabolic states based on pathway membership, phenotypic directionality, and clinical consistency, with broad representation across lipid, carbohydrate, amino-acid, and cofactor/vitamin metabolism. Notably, more than 70,000 signatures aligned with regulated cell-death processes—especially autophagy-dependent cell death and ferroptosis—underscoring the tight coupling between metabolic reprogramming and RCD. Tumor stemness accounted for most significant phenotypic associations, with TMB and MSI contributing secondarily, while immune-cold patterns predominated across metabolic categories. Clinically, signatures segregated into protective and high-risk groups, reflecting divergent metabolic fitness states. Altogether, this atlas offers a structured overview of metabolic behavior across cancers and forms the analytical foundation for reconstructing metabolic regulatory circuitries.

By grouping elements only when they share the same metabolic pathway, RCD mechanism, omic–phenotypic concordance, and immune microenvironment type, over 240,000 metabolic signatures specific to each omic layer were generated, in which multimolecular signatures represent phenotypes coordinated by multiple components, while single-component signatures indicate isolated or context-specific dependencies. Because many metabolic enzymes and regulatory RNAs take part in different pathways and death mechanisms, we allowed the same molecule to appear in multiple signatures, preserving its pleiotropy. This redundancy is therefore biological, not methodological.

The metabolic signatures showed strong associations with the tumor stem-like phenotype (TSM). The recurrent presence of this type of relationship across several omic layers shows that metabolic reprogramming is a defining feature of tumor states with stem-cell-like properties, in which metabolism acts as a central axis of phenotypic plasticity.

For example, serine metabolism reprogramming shows that increased synthesis and utilization favor biosynthetic production, redox control, and chromatin modification, supporting oncogenesis, stemness, and therapeutic resistance. ^[23]^ Similarly, inosine-induced reprogramming in engineered immune cells improves mitochondrial function, polyamine synthesis, and epigenetic remodeling toward a stem-like state, increasing persistence and efficacy of CAR-T cells. ^[24]^ Furthermore, the dependence of several tumor and immune populations on fatty acid β-oxidation (FAO) for survival, maintenance of stemness, metastatic potential, and immune evasion reinforces the central role of metabolic adaptability in determining cell fate under the pressures of the tumor microenvironment. ^[25]^ T-cell stemness itself is metabolically constrained in this microenvironment, where high levels of extracellular K⁺, lactate, and acidosis modulate the balance between self-renewal and terminal exhaustion, affecting the effectiveness and durability of immunotherapeutic responses. ^[26]^ In summary, tumor metabolism plays a central role in phenotypic plasticity, and stemness emerges as a metabolically sustained cellular condition regulated across multiple omic layers.

### 3.3. Convergent and Divergent Metabolic Regulatory Circuitries

A central question emerging from the signature compendium is whether the molecules within each signature—especially multi-member configurations—are connected by common upstream regulatory elements. To address this, we systematically intersected experimentally validated miRNA–mRNA and lncRNA–mRNA interactions across all components of every signature. For enzyme-gene signatures, only regulators that interact with all signature members were retained; for miRNA- and lncRNA-based signatures, the criterion was reversed, identifying coding genes co-regulated by all non-coding elements. This approach ensures that every retained regulator reflects a complete cross-connectivity pattern within the signature rather than partial or incidental interactions.

This analysis revealed extensive regulatory integration across cancer types: over 75% of all metabolic signatures exhibited at least one shared regulatory interaction, providing a structural substrate for reconstructing convergent and divergent regulatory circuitries. Multi-element signatures frequently mapped to multiple shared regulators, indicating points of coordinated metabolic control, whereas single-element signatures often carried broad regulatory connectivity reflecting the pleiotropy of individual metabolic genes and ncRNAs.

A representative example occurs in PAAD, where the metabolic signature PAAD-1340.5.4.0.1.6.3.N.3.3.0.3.3, composed of *NNT* and *SIRT4*—both annotated to nicotinate and nicotinamide metabolism within the cofactors-and-vitamins category—shared five upstream miRNAs: hsa-miR-103a-3p, hsa-miR-15a-5p, hsa-miR-15b-5p, hsa-miR-16-5p, and hsa-miR-195-5p. The presence of this multi-miRNA regulatory module illustrates how functionally coordinated metabolic units can arise from the intersection of biochemical relatedness with common post-transcriptional control.

Together, these findings demonstrate that metabolic signatures are not merely co-annotated metabolic units but regulatory modules structurally anchored in shared ncRNA control, thereby enabling the definition of higher-order metabolic regulatory circuitries.

Another important finding was the predominance of cold immune phenotypes across different tumor types, reinforcing the idea that metabolic reprogramming is closely linked to immune cell suppression and exclusion. This pattern is consistent with evidence in non–small cell lung cancer, where metabolic alterations and suppressive stromal networks sustain inactive immune microenvironments and promote resistance to checkpoint blockade therapy. ^[27]^ Similarly, in breast cancer, MIF-mediated glycolysis not only stimulates tumor growth and dissemination but also inhibits cytotoxic immunity; its inhibition restores pro-inflammatory immune infiltration and increases the effectiveness of checkpoint inhibitors. ^[28]^ In addition, studies show that early immunosurveillance itself can direct the metabolic reprogramming of tumor cells through non-canonical IFNγ–STAT3 signaling, shaping bioenergetic states that allow cancer cells to compete with infiltrating lymphocytes and maintain an immunosuppressive environment. ^[29]^ Taken together, these observations indicate that most signatures classified as indicative of cold immune profiles reflect immune-evasion strategies dependent on tumor metabolism.

A representative example is the lipid metabolism–related signature composed of the genes *AKR1C1* and *AKR1C2*, involved in steroid hormone biosynthesis and the ferroptosis pathway. This signature showed concordance across different omic layers, as its association with the stemness phenotype was observed in both gene expression and CpG methylation profiles, encompassing multiple tumor types, including LUAD, GBM, KIRP, PAAD, PRAD, SKCM, THYM, UCS, and KIRC. In all these contexts, it maintained a consistent correlation with cold immune microenvironments, reinforcing the notion that metabolic alterations associated with stemness often coincide with suppressed or desert immune profiles.

In prostate cancer, the signature involving *AKR1C1* and *AKR1C2* (PRAD-1119.4.13.6.0.6.3.N.2.5.0.1.3) showed an adverse prognostic effect, evidenced by Cox regression analyses for overall survival (p = 0.0136) and disease-specific survival (p = 0.00318). This result is consistent with the known role of AKR enzymes in androgen metabolism. AKR1C2, for example, converts dihydrotestosterone (DHT) into its less active metabolite, 3α-diol; reduced activity of this enzyme in prostate cancer leads to excessive retention of DHT and favors tumor progression. Pharmacological interventions targeting AKR1C2 have been shown to reduce tumor growth both in vitro and in vivo. ^[30]^ In parallel, AKR1C3 enhances androgen biosynthesis, promoting castration resistance and establishing itself as a validated therapeutic target. ^[31,32]^ AKR1C1 also plays an important role in hormonal metabolism, particularly in regulating progesterone homeostasis, and is highly expressed in various hormone-dependent neoplasms. Recent structural and biochemical studies have confirmed the feasibility of developing selective AKR1C1 inhibitors — including synthetic and natural compounds — based on differences in functional domains, co-crystallized binding sites, and polymorphic variants that modulate enzymatic activity and disease susceptibility. ^[33]^

Thus, the recurrence of the *AKR1C1*/*AKR1C2* signature across different tumor types, its association with transcriptional and epigenetic states related to stemness, and its consistent relationship with cold immune microenvironments illustrate how metabolism, particularly lipid metabolism, simultaneously shapes tumor characteristics and immune-evasion mechanisms. This example shows the functional value of representing metabolic dependencies as modular and context-specific signatures, capable of more accurately reflecting the complexity of interactions between metabolism, tumor phenotype, and immune response — in contrast to approaches that reduce these phenomena to one-dimensional classifications.

Among the 241,415 metabolic signatures constructed, 23.6% contained multiple components, whereas 76.4% comprised a single molecule. In the group of multimolecular signatures, 19.3% shared at least one common regulator, although most did not. In contrast, among single-component signatures, 92.8% were associated with at least one regulator, and only a small fraction had none.

An example of a multimolecular signature with shared regulators was identified in PAAD. This signature, composed of the genes *NNT* and *SIRT4* (PAAD-1340.5.4.0.1.6.3.N.3.3.0.3.3), is related to cofactor and vitamin metabolism, specifically nicotinate and nicotinamide metabolism. Both share five regulatory microRNAs — hsa-miR-103a-3p, hsa-miR-15a-5p, hsa-miR-15b-5p, hsa-miR-16-5p, and hsa-miR-195-5p — illustrating how multiple molecules can converge under the control of the same set of regulators.

Based on this analysis, we sought to understand the regulatory relationships between molecules and metabolic signatures that may exhibit convergent or divergent patterns across different biological layers within the same tumor type.

Among 204,591 regulator–signature interactions, divergent patterns predominated (27.2%) compared with convergent ones (24.4%). The immune infiltration domain showed the highest proportion of significant interactions (48.4%), indicating that the immune context represents the main axis of regulatory reconfiguration between signatures and their controllers. Because a single regulator can exert pleiotropic action on several target genes in distinct metabolic pathways, a signature may exhibit both convergent and divergent interactions. This means that the observed effects may result not only from the direct interaction with the signature but also from indirect influences on other targets that differentially modulate the immune, phenotypic, or prognostic landscape of the tumor.

An example of a complex regulatory pattern was observed in the microRNA-based signature (hsa-miR-512-3p + hsa-miR-520e), identified in lung adenocarcinoma (LUAD-1716.2.13.0.1.4.1.P.2.3.3.2.3). This signature is associated with carbohydrate metabolism, particularly the pyruvate pathway, and is linked to the LDHD gene. While the signature itself showed a positive correlation with TMB (ρ = 0.188), LDHD exhibited a negative correlation with TMB (ρ = –0.180). The opposing directions of these correlations indicate a divergent association between the signature and its regulatory partner. Although both microRNAs belonged to the cold immune class, they maintained convergent immune concordance. Survival analyses, however, showed contrasting results: the signature exhibited a risk effect for progression-free interval (PFI), whereas the interaction was protective, generating divergence between Cox regression models and survival analyses. In summary, this signature presented immune convergence but phenotypic, Cox, and survival divergence — reflecting the complexity of regulatory interactions that support phenotypic variability in cancer.

Another example was observed in BLCA, with the transcript-level signature (ENST00000367321 + ENST00000618312) (BLCA-8435.5.5.0.0.5.3.P.3.8.8.1.2), associated with cofactor and vitamin metabolism, particularly the one-carbon pool by folate pathway, which is essential for nucleotide synthesis, methylation, and redox control. This signature interacted with microRNAs hsa-miR-24-3p and hsa-miR-29b-2-5p, both related to TSM phenotypes. Despite positive and convergent correlations, the biological implications of each were distinct: miR-24-3p appeared in a hot immune context but showed divergent immune concordance, whereas miR-29b-2-5p, classified as variable, displayed convergent immune concordance. In survival analyses, miR-24-3p showed a convergent risk effect in OS, DSS, and PFI, whereas miR-29b-2-5p exhibited a protective effect, evidencing a decoupling between transcriptional regulation and phenotypic behavior. Thus, while miR-24-3p combined immune divergence with phenotypic and Cox convergence, miR-29b-2-5p presented immune and phenotypic convergence but divergence in Cox and survival outcomes.

An illustrative example of this divergence is observed in LGG, where the lipid metabolism signature composed of *FADS1* and *SCD* (LGG-1494.4.2.6.1.6.3.P.3.8.8.2.3)—involved in unsaturated fatty acid biosynthesis and ferroptosis—displayed a distinct regulatory pattern in relation to microRNAs miR-215-5p and miR-155-5p. This signature showed a positive correlation with the tumor stemness index (ρ = 0.216), was associated with a cold immune microenvironment, and showed a protective effect in Cox analysis. In contrast, the microRNAs showed negative correlations with the stemness index (ρ = –0.529), were associated with a hot immune phenotype, and were predictors of worse prognosis.

From a functional perspective, *FADS1* and *SCD* exert complementary yet antagonistic roles in regulating lipid metabolism. FADS1 catalyzes the synthesis of polyunsaturated fatty acids, promoting lipid peroxidation and increasing susceptibility to ferroptosis. In triple-negative breast cancer, for example, coexpression of *FADS1* and *FADS2* is associated with poor prognosis, and their pharmacological inhibition reduces both peroxidation and ferroptotic cell death. ^[34]^ In colorectal cancer, FADS1 takes part in an axis that stimulates tumor growth. ^[35]^ Furthermore, the transcription factor Zeb1 activates FADS1 and lipid cofactors such as ACSL4 and ELOVL5, linking mesenchymal plasticity to a lipid-dependent ferroptotic program. ^[36]^

On the other hand, SCD converts PUFAs into monounsaturated fatty acids (MUFAs), reducing lipid peroxidation and conferring resistance to ferroptosis. In gastric cancer, SCD is stabilized by USP7-mediated deubiquitination, and its inhibition by the compound DHPO restores ferroptotic sensitivity. ^[37]^ In triple-negative breast cancer, overexpression of SCD1/2 maintains resistance to ferroptosis, requiring simultaneous inhibition of FADS1/FADS2 for reversal. ^[34]^ In pancreatic adenocarcinoma, SCD sustains cell survival under lipid restriction, and its inhibition reduces tumor viability. ^[38]^ Interestingly, Zeb1-mediated repression of SCD restores vulnerability to ferroptosis, establishing a functional link between lipid metabolism and cellular plasticity. ^[36]^

In the LGG context, the protective effect of the *FADS1*+*SCD* signature, even in the presence of a cold immune microenvironment, suggests that the ratio between PUFAs and MUFAs creates a balanced redox state capable of maintaining a basal ferroptotic vulnerability. While FADS1 promotes ferroptosis sensitivity, SCD acts as a partial compensatory mechanism, preventing excessive oxidative damage. In slow-growing tumors with low inflammation, this metabolic balance may limit both tumor progression and immunopathological damage, in agreement with previous observations regarding these enzymes. ^[34,38]^

Regarding shared interactions of *FADS1*+*SCD*, in the immune system, both miR-215-5p and miR-155-5p act as potent activators of the immune response, promoting CD8⁺ lymphocyte cytotoxicity through the induction of IFNγ and granzyme B, especially when transferred via extracellular vesicles from CD4⁺ T cells. ^[39]^ Their association with a hot immune phenotype but an unfavorable prognosis in LGG indicates a strongly context-dependent effect, possibly related to reactive infiltration in more aggressive tumors, functional differences between tumor and immune cells, and the influence of genomic covariates such as IDH1/2 mutations, 1p/19q co-deletion, and TERT mutations, which reshape the interface between metabolism and immunity. Thus, although their immunostimulatory functions are consistent with the literature, ^[39]^ their clinical impact in LGG is determined by the molecular and cellular landscape of the tumor.

From a therapeutic perspective, co-exploration of metabolic and immunological targets emerges as a promising strategy. Inhibition of *SCD* or *FADS1*/*FADS2*, combined with ferroptosis induction, may potentiate immune responses mediated by miR-215-5p and miR-155-5p. ^[34,36–39]^ In this way, the FADS1+SCD signature may act as a biomarker of ferroptosis sensitivity, while specific miRNA profiles may indicate reactive inflammatory states useful for guiding immunotherapy combinations. As evidenced in studies in colorectal cancer, ^[35]^ monitoring FADS1-dependent eicosanoid signaling is crucial to prevent pro-tumorigenic side effects, reinforcing the need for contextually calibrated therapeutic approaches that integrate lipid metabolism, ferroptosis, and tumor immunity.

In chromophobe renal cell carcinoma (KICH), the metabolic signature composed of *ACAT2* and *HADH*, regulated by miR-124-3p and involved in the degradation of valine, leucine, and isoleucine, in tryptophan metabolism, and in the butanoate pathway, showed a negative correlation with TSM (ρ = –0.52 and –0.45), being associated with a cold immune microenvironment and poor prognosis.

Previous evidence shows that miR-124-3p acts as a tumor suppressor, repressing oncogenic genes such as *EZH2* in prostate cancer, *HRCT1* in gastric cancer, and *RhoG* in glioblastoma, inhibiting proliferation and motility. ^[40,41]^ In KICH, the simultaneous reduction of miR-124-3p and its targets may reflect a compensatory mechanism that limits metabolic stress within an immunosuppressive environment.

The oncogenic enzyme ACAT2 promotes a glycolytic–epigenetic circuit by inducing histone lactylation (H3K18la) and acetylation of the mitochondrial carrier MTCH2, which increases lactate secretion and stimulates M2 macrophage polarization. ^[42]^ The association of *ACAT2*+*HADH* with a cold immune phenotype reinforces the role of *ACAT2* in immunometabolic remodeling and immune evasion. *HADH*, in turn, exhibits a dual and context-dependent behavior: it acts as a tumor suppressor in hepatocellular, gastric, and clear cell renal carcinomas, but as an oncogene in colon cancer and acute myeloid leukemia. ^[43,44]^ In renal tumors, reduced *HADH* correlates with AKT pathway activation, PTEN loss, worse clinical outcome, low CD8⁺ lymphocyte infiltration, and increased Tregs. Thus, the combined metabolic repression of *ACAT2* and *HADH* characterizes a mitochondrial–immunosuppressive phenotype in which impaired β-oxidation and altered cholesterol metabolism reshape the tumor immune landscape.

Taken together, these findings reveal an integrated miR-124-3p/ACAT2/HADH axis that connects epigenetic regulation, energetic metabolism, and immune evasion in KICH. The coordinated reduction of this axis defines tumors metabolically adapted to immunosuppression, suggesting that restoring miR-124-3p expression or rebalancing the ACAT2–HADH pair may reverse immunometabolic resistance and enhance the response to immunotherapy.

In PAAD, a convergent multi-omic regulatory circuitry was identified, connecting fatty acid metabolism to miRNA regulation. This network integrates a transcriptional signature associated with fatty acid degradation—composed of the *ALDH7A1* transcripts ENST00000503281 and ENST00000553117—and a non-coding RNA regulatory signature formed by hsa-let-7b-5p and hsa-miR-193b-3p. Both showed negative correlations with the MSI phenotype and were associated with poor prognosis in OS, DSS, and DFI. They presented a cold immune pattern, characterized by low effector lymphocyte infiltration, outlining a transcriptional–post-transcriptional module that sustains aggressive and immunosuppressed tumors.

The gene *ALDH7A1* is highly expressed in pancreatic cancer cell lines and is related to unfavorable outcomes and a strong dependence on fatty acid oxidation (FAO). ^[45]^ Its function involves detoxifying lipid aldehydes under oxidative stress, generating NADH, and connecting redox balance to energy production. Inhibition of *ALDH7A1* reduces oxygen and ATP consumption, leading to tumor regression in preclinical models. ^[45]^ The observed pattern of high *ALDH7A1* expression combined with low MSI and a cold immune microenvironment reinforces a model in which the activity of this enzyme creates a redox–energetic state that favors tumor growth and hampers the antitumor immune response.

This behavior is consistent with studies in non–small cell lung cancer and breast cancer, which associate *ALDH7A1* with so-called “immune-desert” microenvironments. ^[46]^ Furthermore, enzymes of the ALDH family, including ALDH7A1, sustain cancer stem cell programs, promoting self-renewal, drug resistance, and tumor progression. ^[47]^ *ALDH7A1* overexpression also activates the JAK–STAT and mTOR pathways, contributing to the maintenance of cell growth. ^[48]^ Thus, the association between poor prognosis and suppressed immunity likely reflects the presence of metabolically adapted tumor subpopulations capable of surviving oxidative stress and immune pressure.

From a regulatory perspective, hsa-let-7b-5p is a tumor suppressor miRNA known for restricting proliferation and mitotic escape by targeting genes such as *TK1*, *AURKB*, *BIRC5*, *PLK1*, and *EZH2*, promoting apoptosis and increased cell survival. ^[42,49–52]^ Loss of its function leads to derepression of metabolic and proliferative pathways. hsa-miR-193b-3p acts similarly, modulating genes involved in FAO and apoptosis. The functional convergence between let-7b-5p and miR-193b-3p suggests the existence of a cooperative metabolic control module whose dysregulation amplifies oxidative metabolism and reinforces immune exclusion in PAAD.

Taken together, these results describe a metabolic–immunosuppressive circuitry in which ALDH7A1 drives oxidative metabolism and immune evasion, while the loss of let-7b-5p and miR-193b-3p removes the post-transcriptional brakes that would otherwise contain this process. This integration between metabolic reprogramming, immunosuppression, and adverse clinical outcome positions oxidative metabolism as a central element of tumor aggressiveness in pancreatic cancer.

Our results support a model in which tumor metabolism functions as an n-dimensional regulatory space, interdependently integrating genetic, epigenetic, and transcriptomic layers with tumor, immune, and clinical phenotypes. Within this analytical framework, a metabolic regulatory circuitry is defined as a paired relationship between two omic signatures in a specific cancer type: one composed of non-coding RNA regulators and the other formed by metabolically validated targets. Each regulatory element interacts with each target element, forming coherent pairs whose combined behavior determines whether control is convergent or divergent across molecular, phenotypic, and clinical layers. Divergent circuitries reflect the canonical regulatory logic, in which regulator and target exert opposite biological effects — for example, one associated with risk and the other with protection. Conversely, convergent circuitry shows cooperative reinforcement, in which both act in the same direction on metabolism, phenotype, and prognosis. Together, these modules outline the architectures through which non-coding RNAs modulate metabolic state, tumor progression, and the immune context.

### 3.4. Strengths and limitations

The strength of this work lies precisely in its integrative nature, conceptual innovation, and operational applicability. The study proposes an analytical framework capable of unifying multiple omic layers within an n-dimensional regulatory space, in which metabolic signatures and their non-coding regulators are treated as complex informational entities rather than mere gene lists. This multidimensional perspective transcends traditional linear analysis, enabling signatures to be represented and compared by their geometry — in terms of volume, coherence, and orientation — and thus distinguishing mechanistic convergence from simple redundancy. This geometric formalization not only enhances the understanding of the regulatory architecture of tumor metabolism but also establishes the foundations for a new mode of thinking about biological interactions in topological and relational terms.

In this context, OncoMetabolismGPS emerges as a practical translation tool for this concept. Developed in Shiny, the application offers an interactive interface that integrates molecular composition, tumor phenotypes, immune states, and prognostic data in a unified exploratory environment. By enabling dynamic navigation across omic and phenotypic layers, the system expands discovery potential, making it possible to explore regulatory circuitries from complex public datasets.

From a translational perspective, the metabolic signatures identified show potential to improve patient stratification, guide targeted therapies, and reveal specific metabolic vulnerabilities. The fact that different signatures share common regulators suggests such molecules may act as nodal control points whose modulation could shift tumors between metabolic states associated with aggressiveness, immune exclusion, or susceptibility to RCD. In one of our signatures, this principle — exemplified by the pathways mediated by FADS1/FADS2/SCD, USP7–SCD, and enzymes such as HADH and ACAT2 — illustrates the power of the framework in coherently and biologically meaningfully integrating functional and phenotypic information. ^[34,35,37,39,40,43,44,53,54]^

Beyond its translational value, the conceptual aspect developed in this study opens new possibilities for basic science. The geometric representation of signatures and their interactions enable the exploration of biological system organization from a mathematical and topological perspective, offering an alternative path to investigate functional coherence, adaptive redundancy, and regulatory modularity. Such an approach may foster the formulation of mechanistic hypotheses, reveal emerging principles of molecular organization, and contribute to the discovery of new biological mechanisms. Thus, the proposed framework not only provides tools for identifying biomarkers and therapeutic targets but also establishes a conceptual proposal for advancing systems biology.

Despite its conceptual and methodological advances, this study presents limitations that must be acknowledged in a balanced manner. The inferred regulatory interactions were based on multi-omic correlations and alignment across molecular, phenotypic, and clinical domains, which enables the formulation of mechanistic hypotheses but not the establishment of direct causal relationships. Although powerful for uncovering emerging patterns, this approach still relies on complementary experimental validations to confirm the proposed functional links. The analyses were conducted predominantly using bulk RNA-Seq data, which may dilute the contribution of cellular subpopulations and mask metabolic states specific to certain cell types or microenvironmental niches. The future incorporation of spatial and single-cell transcriptomics data may substantially refine the inferences presented here, allowing the identification of regulatory circuitries that are contextually restricted. Another point to consider is that the annotation of metabolic pathways and modules depends on public databases such as KEGG, which may not fully reflect context-specific metabolic reprogramming or evolutionary adaptations in different tumor environments. Finally, although the convergence–divergence framework provides a solid theoretical basis for integrating metabolism, tumor phenotype, immune state, and prognosis, the therapeutic implications derived from these relationships remain exploratory and require more extensive functional and clinical testing.

## 4. Conclusion

This work introduces a unified, multi-omic framework that resolves tumor metabolism into modular, context-specific signatures and their shared regulatory circuitries, linking metabolic pathways to RCD mechanisms, tumor phenotypes (MSI, TMB, TSM), immune ecologies (hot/cold/variable), and clinical outcomes across 33 cancers. By systematically integrating enzyme-coding genes, non-coding regulators (miRNAs, lncRNAs), and transcript isoforms, and by enforcing concordance across molecular, phenotypic, immune, and prognostic dimensions, we delineate a layered regulatory architecture in which biochemical activity, cell-fate programs, and microenvironmental pressures co-evolve. The resulting atlas of >240k omic-specific metabolic signatures reveals both conserved and tumor-type-restricted dependencies, with a predominance of immune-cold states and extensive domain-specific convergence/divergence between signatures and their regulators -an organizational principle rarely interrogated in prior studies.

Two mechanistic patterns emerge with translational relevance. First, convergent interactions (e.g., *ACAT2* + *HADH* with miR-124-3p) denote stable disease-driving programs in which metabolic dependency and regulatory control align across molecular, immune, and clinical domains. Second, divergent interactions (e.g., *FADS1* + *SCD* with miR-155-5p/miR-215-5p) expose adaptive fault lines where proliferative/stemness advantages trade off against immune visibility or ferroptosis liability, suggesting entry points for therapeutic sensitization. Together, these patterns argue that metabolic state is not a single hallmark but an adaptive, reconfigurable regulatory landscape whose topology can be read out by signatures and perturbed through shared regulators.

Beyond cataloging, OncoMetabolismGPS operationalizes this framework, enabling hypothesis generation, patient stratification by metabolic state, prioritization of regulators as sensitizers, and rational design of combination strategies that pair metabolic rewiring with cell death-linked interventions and immunotherapy. While causal directionality and cell-type resolution require experimental perturbation and single-cell/spatial validation, the present study establishes the conceptual and analytical foundations for prospective biomarker development and biomarker-guided trials.

## 4. Experimental Section

### 4.1. Identification of metabolism-related coding and non-coding RNAs

We retrieved 1,406 human enzyme-coding genes involved in anabolic and catabolic reactions across 127 KEGG-defined metabolic pathways, grouped into seven major metabolic categories: carbohydrate metabolism, lipid metabolism, energy metabolism, amino acid metabolism, metabolism of other amino acids, nucleotide metabolism, and metabolism of cofactors and vitamins. Gene selection was based on the Kyoto Encyclopedia of Genes and Genomes using the KEGGREST package ^[17,55]^.

For these enzyme-coding genes, we identified 2,490 miRNAs and 242 lncRNAs with experimentally validated regulatory interactions. miRNA-mRNA interactions were obtained from miRTarBase ^[56]^, and lncRNA-mRNA interactions from EVlncRNAs 3.0 ^[57]^. For each gene and regulatory RNA, transcript isoform identifiers were retrieved from Ensembl (GRCh38.p13) ^[58]^, totaling 20,592 annotated isoforms.

### 4.2. Annotation of regulated metabolic cell death mechanisms

To determine the involvement of molecular targets in RCD mechanisms with metabolic dependency, we performed functional annotation using a manually curated knowledge base of RCD (RCD) pathways. ^[59]^ Targets were classified according to their participation in ferroptosis, apoptosis, necroptosis, parthanatos, and other metabolic cell death programs.

### 4.3. Multi-omic, phenotypic, and clinical integration across 33 cancer types

To further characterize the molecular targets, we integrated multi-omic, phenotypic, and clinical data from The Cancer Genome Atlas (TCGA) via UCSC Xena, covering 33 distinct cancer types. ^[60]^ The omic layers analyzed included bulk RNA sequencing, CpG methylation, CNV, somatic mutations, and protein expression (RPPA array). In parallel, we incorporated clinical and phenotypic data, including immunogenomic features with potential impact on immunotherapy response, such as microsatellite instability (MSI), tumor mutational burden (TMB), and stemness indices reflecting tumor plasticity and self-renewal capacity, which have been linked to immune evasion, cold tumor microenvironments, and reduced immunotherapeutic efficacy.

### 4.4. Computational pipeline for integrative association testing

We developed an R-based pipeline to systematically evaluate associations between molecular features and tumor phenotypes. Approximately 2.94 million pairwise association tests were performed across cancer types and omic layers.

Association analyses were stratified by cancer type and omic layer. Multiple-testing correction was applied using the Bonferroni method, and associations with adjusted p-values < 0.05 were significant.

### 4.5. Prognostic evaluation of significant associations

Molecular features significantly associated with tumor phenotypes were subsequently evaluated for prognostic value only in the same tumor type and omic layer where the association had been detected. For example, if the expression of a gene was significantly correlated with increased stemness in lung adenocarcinoma (LUAD), its prognostic evaluation was restricted to that specific tumor-omic context.

For each selected instance, we applied univariate Cox proportional hazards regression to estimate the hazard ratio of clinical outcomes, complemented by log-rank tests to compare survival distributions between stratified groups in clinical endpoints, including OS, DSS, DFI, and PFI.

### 4.6. Characterization of the tumor immune microenvironment

To characterize the tumor microenvironment linked to the omic layers of significant targets, we analyzed correlations between molecular features and infiltrating immune cell populations. Relative abundance estimates for 29 immune cell types were derived using the CIBERSORT and xCELL deconvolution method applied to TCGA transcriptomic profiles (Table S8). ^[61,62]^ Each cell type was categorized into anti-tumor, pro-tumor, or dual functional groups based on literature evidence.

Omic layers were then classified according to the predominant associated immune infiltration pattern. For refined categorization, we focused on five immune cell subsets -CD8⁺ T cells, NK cells, regulatory T cells (Tregs), M1 macrophages, and M2 macrophages -widely recognized as markers of hot versus cold tumors. Associations dominated by effector infiltrates (CD8⁺, NK, M1) were classified as hot, those dominated by suppressive infiltrates (Tregs, M2) as cold, and intermediate or mixed profiles as variable (Supporting Information – Methods A and B).

### 4.7. Construction of omics-specific metabolic signatures

To integrate multi-omic associations into biologically interpretable units, we constructed omic-specific metabolic signatures for each molecular class (mRNAs, miRNAs, lncRNAs, and transcript isoforms). A metabolic signature was defined as a set of molecules that jointly satisfied a set of four hierarchical criteria, ensuring both mechanistic coherence and translational relevance. Signature construction was performed independently within each cancer type to preserve tumor-context specificity.

First, candidate elements had to share functional grounding in metabolism, evidenced by (i) participation in the same KEGG metabolic pathway and (ii) annotation to the same metabolism-RCD mechanism (RCD database). Second, all elements in a signature were required to exhibit concordant associations with tumor phenotypes (MSI, TMB, and/or stemness indices) within a given tumor type and omic layer. Third, these associations needed to display consistent prognostic directionality across survival outcomes (OS, DSS, DFI, PFI), as determined by Cox proportional hazards models and validated by log-rank testing. Finally, signatures were required to exhibit immunological compatibility, meaning that all components were embedded in tumor microenvironments characterized by the same immune state (hot, cold, or variable), as inferred from immune infiltration profiles.

Elements that met all four criteria were grouped into multi-component signatures when they shared identical biological and clinical association profiles across dimensions. When a candidate element had no additional partners meeting these criteria, it was kept as a single-component signature, preserving isolated yet potentially critical metabolic features.

Because some metabolic genes and their regulatory RNAs participate in multiple pathways and cell death processes, the same molecule could appear in more than one signature. This controlled redundancy is a deliberate feature of the framework: it captures the multifunctionality and pleiotropic roles of metabolic regulators rather than collapsing them into a single assignment. By allowing biological context to guide grouping rather than forcing mutual exclusivity, the resulting signatures more accurately reflect the true network architecture of tumor metabolism.

### 4.8. Identification of shared regulatory interactions

To determine common regulatory control, experimentally validated miRNA-mRNA and lncRNA-mRNA interaction tables were intersected across all members of each signature. A regulator was kept only if it interacted with every component of the signature. The same logic was applied inversely for miRNA-based signatures to identify shared coding targets. The algorithm intersected the interactions of all components within each signature, keeping only regulatory molecules or targets common to all members. For example, in a signature composed of two enzyme-coding genes initially linked to miR-21, miR-155, and miR-35, only miR-21 interacted with both genes and was therefore recorded as the validated shared regulator. Similarly, for miRNA-based signatures, shared coding gene targets were identified.

### 4.9. Convergence versus divergence analysis of regulatory circuitries

For each cancer type, we evaluated whether the signature and its shared regulator showed aligned (convergent) or opposing (divergent) associations with tumor phenotypes (MSI, TMB, stemness), survival outcomes (Cox and log-rank analyses), and immune microenvironment classifications.

Interactions were convergent when both the signature and its shared regulator showed the same direction of association (e.g., both protective in OS or both enriched in anti-tumor microenvironments). Conversely, they were classified as divergent when associations were opposite, highlighting potential compensatory mechanisms in tumor biology.

### 4.10. Signature nomenclature system

We designed a standardized signatuwre nomenclature system to ensure traceability and clarity across computational analyses and visual outputs. Every signature was assigned a unique alphanumeric identifier incorporating: (i) tumor type, (ii) omic layer, (iii) metabolic pathway category, (iv) metabolic cell death involvement, (v) presence of shared regulatory interactions, (vi) phenotypic association class (MSI, TMB, TSM), (vii) prognostic classification across clinical outcomes, and (viii) tumor microenvironment and immune infiltration profiles. A full description of all encoding rules, scoring weights, and rank calculations is provided in the Supporting Information (Method C).

## Supporting Information

Supporting information is available from the journal online library.

## Acknowledgements

The results shown here are based on data and resources generated by the TCGA Research Network (https://www.cancer.gov/tcga), UCSC Xena (https://xena.ucsc.edu), UCSC Xena Shiny (https://shixiangwang.shinyapps.io/ucscxenashiny/), and KEGG (https://www.kegg.jp).

## Author contributions

H.A.C.N. conceptualized the study; developed, validated, and implemented the R-based computational pipeline; performed integrative analyses; generated the initial draft; contributed to manuscript revision; and developed the Shiny application and GitHub repository.

E.R.S. contributed to pipeline development and code validation; generated figures; and participated in manuscript editing.

V.S.L. generated figures and contributed to manuscript editing.

E.M.A. conceptualized and supervised the study; contributed to methodological design; developed, validated, and audited the computational pipeline; and contributed to writing and critical revision of the manuscript.

## Data Availability Statement

The data and source codes that support the findings of this are openly available at the GitHub URL: https://github.com/HigorACNogueira/Multi-omic-Oncometabolism-GPS

