## Supporting information for "A Multi-Omic Atlas of Convergent and Divergent Metabolic Regulatory Circuitries in Cancer"

H.A.C.N. Author 1, E.R.S. Author 2, V.S.L. Author 3, E.M.A. Author 4

Laboratório de Biotecnologia, Centro de Biociências e Biotecnologia, Universidade Estadual do Norte Fluminense, Campos dos Goytacazes, Brazil.


### **Supporting information - Methods**

### **A. Microenvironment classification (anti-tumoral / pro-tumoral / dual)**

### This analysis assessed how each molecular feature relates to the broader immune context of the tumor. Relative abundances of 29 immune cell types were obtained from TCGA bulk transcriptomic data using the CIBERSORT and xCELL algorithms. Based on literature-supported functional roles, these cell types were grouped into anti-tumoral, pro-tumoral, or dual / context-dependent categories (Supplementary Table S8).

##### **A.2 Correlation interpretation**

Spearman correlations were computed independently for each cancer type, and only correlations with p < 0.05 were kept. For transcriptomic features with an assigned expression state, the sign of the correlation was interpreted conditionally:

- When a feature was overexpressed, a positive correlation (ρ > 0) was interpreted as an increase in the infiltrate (direct), whereas a negative correlation (ρ < 0) was interpreted as a reduction in the infiltrate (inverse).
- When a feature was underexpressed, a positive correlation showed a reduction in the infiltrate (inverse), while a negative correlation showed an increase (direct).
- For unchanged features, positive correlations were interpreted as direct and negative correlations as inverse, analogous to the overexpressed case.

For non-transcriptomic layers (DNA methylation, CNV, mutation, and RPPA), correlation signs were interpreted directly: ρ > 0 showed a direct association, and ρ < 0 showed an inverse association.

**Simplified step-by-step**

Only correlations with *p* < 0.05 were considered.

| **Expression State** | **ρ > 0** | **ρ < 0** |
| --- | --- | --- |
| **Overexpressed** | increased infiltrate (direct) | reduced infiltrate (inverse) |
| **Underexpressed** | reduced infiltrate (inverse) | increased infiltrate (direct) |
| **Unchanged** | increased infiltrate (direct) | reduced infiltrate (inverse) |

For non-transcriptomic layers:

- **ρ > 0** → direct
- **ρ < 0** → inverse

**A.3 Scoring and classification**

For each feature, all significant correlations contributed to a cumulative score per immune cell class. Direct associations were added to the score, whereas inverse associations were subtracted. The magnitude of the association (|ρ|) determined the strength of the contribution.

The overall immune profile of each feature was then assigned based on the immune class with the highest cumulative score: features with a dominant positive score in immune-effector cell classes were classified as anti-tumoral; those with a dominant positive score in immunosuppressive cell classes were classified as pro-tumoral; and those without a clear dominance were classified as dual. This classification reflects whether the feature is associated with a tumor microenvironment favoring immune activation, immune suppression, or a mixed phenotype.

**Simplified step-by-step**

Each significant correlation contributed to a cumulative score per immune cell class:

- Direct associations added to the score.
- Inverse associations subtracted from the score.
- The magnitude of |ρ| determined contribution strength.

The feature was classified according to the class with the highest cumulative score:

| **Highest Score** | **Profile** |
| --- | --- |
| Anti-tumoral | **Anti-tumoral** |
| Pro-tumoral | **Pro-tumoral** |
| Dual | **Dual** |

This profile reflects whether the feature is associated with a microenvironment that supports immune attack, immune suppression, or mixed behavior.

**B. Immune Phenotype Classification (hot / cold / variable).**

This method focuses specifically on classifying the immune state of the tumor microenvironment. In contrast with Method A, which incorporates 29 immune cell subsets, this classification relies on a reduced set of canonical immune markers that are broadly recognized as indicators of immune activation or suppression. The effector (“hot”) phenotype was defined based on enrichment of CD8⁺ T cells, activated NK cells, M1 macrophages, activated CD4⁺ memory T cells, and activated dendritic cells. Conversely, the suppressive (“cold”) phenotype was defined based on the presence of regulatory T cells (Tregs) and M2 macrophages. Tumor samples exhibiting a mixed or non-dominant pattern across these cell classes were designated as variable, reflecting an intermediate or context-dependent immune microenvironmental state.

- **Effector / Hot markers:**
   CD8⁺ T cells, activated NK cells, macrophages M1, activated CD4⁺ memory T cells, activated dendritic cells
- **Suppressive / cold markers:**
   Regulatory T cells (Tregs), macrophages M2

#### **B.1 Interpretation rules**

Correlation directions were interpreted according to the same expression-state framework described in Method A. Thus, for transcriptomic features, the sign of the correlation was evaluated in the context of the feature's expression state, whereas for non-transcriptomic features the correlation sign was interpreted directly.

**B.2 Phenotype assignment**

#### Features were assigned to an immune phenotype based on the direction of association with effector versus suppressive immune subsets. Specifically, Hot phenotypes were defined by direct associations with effector cell subsets and inverse associations with suppressive subsets. Conversely, cold phenotypes were defined by direct associations with suppressive subsets and inverse associations with effector subsets. Features exhibiting concurrent effector and suppressive association patterns were classified as Variable, reflecting a mixed or context-dependent immune behavior.

- **Hot:** direct association with effector subsets and inverse association with suppressive subsets.
- **Cold:** direct association with suppressive subsets and inverse association with effector subsets.
- **Variable:** co-occurrence of both effector and suppressive associations.

**B.3 Tie resolution**

When both Hot and Cold signals were present at comparable strength, the feature was classified as Variable. If the strongest signal was tied between Hot and Variable, the feature was classified as Hot. Likewise, if the strongest signal was tied between Cold and Variable, the feature was classified as Cold.

- If Hot and Cold signals co-occur → **Variable**
- If Hot ties with Variable → **Hot**
- If Cold ties with Variable → **Cold**

**C. Nomenclature of omic-specific metabolic signatures**

We developed a compact and lossless nomenclature system to (i) uniquely identify each metabolic signature within its tumor type, (ii) encode its metabolic context, omic and phenotypic associations, survival summary patterns, and immune classification outputs, and (iii) preserve machine-readable traceability across all downstream analytical steps and datasets.

**C.1 Identifier structure**

Each signature identifier is represented as a single string composed of one header followed by 13 dot-separated tokens, in the following format:

**[CTAB]-[USI].[MET].[PATH].[MCD].[RCC].[OFC].[PFC].[SCS].[TNC].[HRC].**

**[SMC].[TMC].[TIC]**

The components are defined as follows:

**CTAB:** TCGA tumor code.

**USI:** Unique Series Identifier, a per-tumor sequential integer assigned according to the final post-filtering row order.

**MET:** Metabolism category code corresponding to the broader metabolic domain to which the signature belongs.

**PATH:** Pathway index within the specified metabolic category.

**MCD:** Metabolic cell death mechanism code, showing the RCD pathway with which the signature is associated.

**RCC:** Shared or meaningful interaction flag, showing whether all components share a common regulator interaction.

**OFC:** Omic-layer code specifying whether the signature is composed of genes, miRNAs, lncRNAs, or transcript isoforms.

**PFC:** Phenotypic-layer code representing the associated phenotypic dimension (e.g., stemness, TME infiltration, clinical class).

**SCS:** Spearman correlation direction (P or N) summarizing the dominant regulatory polarity of the signature.

**TNC:** Tumor-versus-normal differential expression code.

**HRC:** Survival hazard-direction template summarizing risk direction across all four clinical endpoints.

**SMC:** Survival worst-group template reflecting consistency of poor-outcome stratification across endpoints.

**TMC:** Tumor microenvironment class as defined in Method A.

**TIC:** Immune phenotype category (Hot, Cold, Variable, or NS), as defined in Method B.

**C.2 Codebooks and Derivation Rules.**

The MET field encodes the metabolic domain to which each signature belongs, following a fixed categorical mapping: 1 = amino acid metabolism, 2 = carbohydrate metabolism, 3 = energy metabolism, 4 = lipid metabolism, 5 = cofactors and vitamins, 6 = other amino acids, and 7 = nucleotide metabolism. Within each metabolic category, distinct KEGG pathways were alphabetically ordered and assigned a PATH index ranging from 1 to K, where K corresponds to the number of pathways within that category; these indices were then mapped back to each signature. The MCD field shows whether the signature is annotated to any metabolic cell-death mechanism (0 = unrelated, 1 = annotated). The INT field denotes a shared or meaningful regulatory interaction among all components of the signature (0 = absent, 1 = present), for example, shared miRNA, lncRNA, or coding-gene regulatory targeting.

Omic-layer and phenotypic-layer encoding is captured through the OFC and PFC fields, respectively. The OFC code identifies the molecular data modality: 1 = protein (RPPA), 2 = mutation, 3 = CNV, 4 = miRNA expression, 5 = transcript (isoform) expression, 6 = gene (mRNA) expression, and 7 = DNA methylation; the numerical order is categorical and does not imply ranking. The PFC code specifies the phenotypic feature associated with the signature: 1 = tumor mutational burden (TMB), 2 = microsatellite instability (MSI), and 3 = stemness. The SCS field records the sign of the dominant Spearman correlation (P = positive, N = negative). The TNC field summarizes tumor-versus-normal expression status using four states: 0 = no corresponding data, 1 = unchanged, 2 = underexpressed, and 3 = overexpressed; these states are later used in immune-profile interpretation.

Two composite templates summarize survival across four clinical endpoints (DSS, DFI, PFI, OS) in a fixed left-to-right order (1–4). The HRC field (“hazard-direction template”) encodes the direction of hazard ratios using letter codes: A = no signal (NS or unavailable), B = risky (HR > 1), and C = protective (HR < 1). For example, 1B2A3C4A shows that the signature is associated with increased risk in DSS, no detectable association in DFI, protective association in PFI, and no detectable association in OS. The SMC field (“survival worst-group template”) uses endpoint-specific letter codes to denote which clinical group exhibited the worst outcome, with interpretation rules dependent on the omic layer. For OFC in {1, 4, 5, 6, 7} (protein, miRNA, transcript, mRNA, methylation), B shows that the high group had worse survival, C shows that the low group had worse survival, and A shows NS/no data. For mutation (OFC = 2), B indicates mutant-worst, C indicates wild-type-worst, and A indicates NS. For CNV (OFC = 3), B indicates deleted-worst, C indicates duplicated/normal-worst, D indicates mixed or compound-worst patterns, and A indicates NS. Signatures with HRC = "1A2A3A4A" and SMC = "1A2A3A4A" were exported as no-effect signatures.

Finally, immune summaries were encoded using the TMC (tumor microenvironment class) and TIC (immune phenotype) fields. TMC was defined as 1 = anti-tumoral, 2 = dual, 3 = pro-tumoral, and 4 = not significant (NS). TIC was defined as 1 = Hot, 2 = Variable, 3 = Cold, and 4 = NS, following the classification rules described in Method B.

**C.3 Example**

The following identifier illustrates the encoding scheme:

BRCA-012.4.7.1.1.6.3.P.3.1B2A3B4A.1B2C3B4A.3.1

In this example, BRCA denotes the tumor type, and 012 is the Unique Series Identifier. The signature belongs to the lipid metabolism category (MET = 4) and corresponds to the 7th pathway within this metabolic class (PATH = 7). The signature is annotated to a metabolic cell death mechanism (MCD = 1) and displays a shared regulatory interaction (INT = 1). The omic layer is gene expression (OFC = 6), and the associated phenotypic axis is stemness (PFC = 3).

The dominant correlation sign is positive (SCS = P), and the signature is overexpressed relative to normal tissue (TNC = 3). The survival hazard-direction template (HRC = 1B2A3B4A) indicates increased risk in DSS and PFI, with no detectable association in DFI or OS. The survival worst-group template (SMC = 1B2C3B4A) reports that the high group had worse DSS and PFI outcomes, the low group had worse DFI outcome, and OS had no significant stratification.

Finally, the tumor microenvironment classification (TMC = 3) indicates a pro-tumoral immune context, and the immune phenotype class (TIC = 1) corresponds to a Hot phenotype.

### **Supporting datasets**

**Dataset S1.**

Matrix of all statistically significant associations (FDR-adjusted p < 0.05) identified across transcriptomic, epigenomic, genetic, and proteomic layers in relation to MSI, TMB, and TSM phenotypes in 33 TCGA tumor types. This dataset provides the foundational association landscape used in subsequent integrative analyses.

**Dataset S2.**

Expanded association resource comprising 463,433 entries generated after annotation to KEGG metabolic pathways, metabolism-linked RCD mechanisms, and experimentally validated regulatory interactions. The dataset captures legitimate multi-mapping arising from the multifunctionality of metabolic enzymes and their non-coding RNA regulators.

**Dataset S3.**

Comprehensive compendium of 241,415 omics-specific metabolic signatures reconstructed across cancer types. Multi-component signatures include elements exhibiting fully concordant molecular, phenotypic, prognostic, and immune profiles; single-component signatures are kept when no additional molecules satisfy the concordance criteria. This dataset defines the metabolic signature landscape used to interrogate regulatory logic.

**Dataset S4.**

Collection of 204,591 regulator–signature interactions derived from the intersection of metabolic signatures with experimentally validated miRNA–mRNA and lncRNA–mRNA networks. Of all signatures, 84,270 exhibit at least one shared regulator. The multi-mapping structure reflects both the pleiotropy of regulatory RNAs and the context-specificity of metabolic programs.

**Dataset S5.**

Atlas of 24,796 metabolic regulatory circuitries obtained by mapping composite interaction signatures from Dataset S4 onto the omic-specific metabolic signatures defined in Dataset S3. Each circuitry is a fully connected regulator–target pair classified as convergent or divergent across molecular, phenotypic, immune, and clinical dimensions, forming the core higher-order regulatory architecture characterized in this study.

### **Supporting information - Figures**

**Figure S1. Pan-Cancer overview of metabolic signatures linked to key molecular tumor phenotypes.** (A) Distribution across omic layers evidencing the proportional contribution of CNV, mutation, methylation, gene, protein, transcript, and miRNA expression data to signatures correlated with microsatellite instability, stemness, and tumor mutational burden. (B) Comparative analysis across metabolic pathways shows a strong enrichment for amino acid, carbohydrate, and energy metabolism. (C) Classification by metabolic cell death uncovers a predominance of apoptosis, expenses, and ferroptosis across tumor types with distinct molecular contexts.

**Figure S2. Pan-Cancer landscape of metabolic signatures according to tumor microenvironment profile**. (A) Layer-wise partitioning highlights the contribution of CNV, mutation, methylation, gene, protein, transcript, and miRNA expression to anti-tumoral, dual, and pro-tumoral profiles. (B) Metabolic stratification reveals consistent representation of amino acid, carbohydrate, and energy metabolism across these biological categories. (C) Distinct cell death modalities—particularly apoptosis, oxieptosis, and ferroptosis—emerge as characteristic mechanisms shaping the metabolic polarity of tumors.

**Figure S3. Immunometabolic distribution of signatures across immune phenotypes.** (A) Relative contribution of CNV, mutation, methylation, gene, protein, transcript, and miRNA expression layers to signatures defining immune microenvironmental states. (B) Functional mapping of metabolic pathways shows dominance of amino acid, carbohydrate, and energy metabolism across immune subtypes. (C) Frequencies of metabolic cell death modes reveal apoptosis, oxieptosis, and ferroptosis as major contributors to the immunometabolic architecture of tumors.

**Figure S4. Pan-Cancer profiles of metabolic signatures associated with overall survival.** (A) Distribution across omic layers showing the representation of each data type (CNV, mutation, methylation, gene, protein, transcript, and miRNA expression) in protective versus risky signatures. (B) Functional classification shows enrichment in amino acid, carbohydrate, and energy metabolism pathways linked to patient outcomes. (C) Distinct death programs—apoptosis, oxieptosis, and ferroptosis—are recurrently observed among survival-related metabolic modules.

**Figure S5. Metabolic signatures related to disease-specific survival in Cox regression across tumor types**. (A) Omic-layer stratification showing the relative weight of CNV, mutation, methylation, gene, protein, transcript, and miRNA expression in protective and risky signatures. (B) Pathway-level decomposition reveals amino acid and energy metabolism as dominant axes in disease-specific metabolic control. (C) Analysis of metabolic cell death modalities identifies apoptosis, oxieptosis, and ferroptosis as central components of prognostic regulation in disease-specific contexts.

**Figure S6. Distribution of metabolic signatures associated with disease-free interval in Cox regression across cancers.** (A) Omic-level contribution analysis for CNV, mutation, methylation, gene, protein, transcript, and miRNA expression data between protective and risky signatures. (B) Comparative enrichment analysis shows consistent participation of amino acid, carbohydrate, and energy metabolism pathways across tumor types. (C) Apoptosis, oxieptosis, and ferroptosis stand out as the most recurrent metabolic death mechanisms influencing recurrence-free survival.

**Figure S7. Pan-Cancer distribution of metabolic signatures associated with progression-free interval in Cox regression.** (A) Contribution of CNV, mutation, methylation, gene, protein, transcript, and miRNA expression layers to protective and risky profiles derived from Cox regression. (B) Metabolic pathway distribution shows sustained dominance of amino acid, carbohydrate, and energy metabolism in progression dynamics. (C) Metabolic cell death characterization highlights apoptosis, oxieptosis, and ferroptosis as the major determinants of tumor progression control across cancer types.
